# Recruitment of BAG2 to DNAJ-PKAc scaffolds promotes cell survival and resistance to drug-induced apoptosis in fibrolamellar carcinoma

**DOI:** 10.1101/2023.06.28.546958

**Authors:** Sophia M Lauer, Mitchell H Omar, Martin G Golkowski, Heidi L Kenerson, Bryan C Pascual, Katherine Forbush, F Donelson Smith, John Gordan, Shao-En Ong, Raymond S Yeung, John D Scott

## Abstract

The DNAJ-PKAc fusion kinase is a defining feature of the adolescent liver cancer fibrolamellar carcinoma (FLC). A single lesion on chromosome 19 generates this mutant kinase by creating a fused gene encoding the chaperonin binding domain of Hsp40 (DNAJ) in frame with the catalytic core of protein kinase A (PKAc). FLC tumors are notoriously resistant to standard chemotherapies. Aberrant kinase activity is assumed to be a contributing factor. Yet recruitment of binding partners, such as the chaperone Hsp70, implies that the scaffolding function of DNAJ-PKAc may also underlie pathogenesis. By combining proximity proteomics with biochemical analyses and photoactivation live-cell imaging we demonstrate that DNAJ-PKAc is not constrained by A-kinase anchoring proteins. Consequently, the fusion kinase phosphorylates a unique array of substrates. One validated DNAJ-PKAc target is the Bcl-2 associated athanogene 2 (BAG2), a co-chaperone recruited to the fusion kinase through association with Hsp70. Immunoblot and immunohistochemical analyses of FLC patient samples correlate increased levels of BAG2 with advanced disease and metastatic recurrences. BAG2 is linked to Bcl-2, an anti-apoptotic factor that delays cell death. Pharmacological approaches tested if the DNAJ-PKAc/Hsp70/BAG2 axis contributes to chemotherapeutic resistance in AML12^DNAJ-PKAc^ hepatocyte cell lines using the DNA damaging agent etoposide and the Bcl-2 inhibitor navitoclax. Wildtype AML12 cells were susceptible to each drug alone and in combination. In contrast, AML12^DNAJ-PKAc^ cells were moderately affected by etoposide, resistant to navitoclax, but markedly susceptible to the drug combination. These studies implicate BAG2 as a biomarker for advanced FLC and a chemotherapeutic resistance factor in DNAJ-PKAc signaling scaffolds.

## INTRODUCTION

Rare tumors represent approximately 20% of total cancer incidence^1^. Gene fusions that involve protein kinases are an emerging class of oncogenes that advance certain rare neuroendocrine and solid tumors^2^. Expression of a DNAJ-PKAc fusion kinase is an emblematic feature of the adolescent liver cancer fibrolamellar carcinoma (FLC)^3^. Approximately 65-100 new cases of FLC are diagnosed in the United States each year, typically in young adults between the ages of 15- 35 with no history of risk factors for liver disease^4^. Five-year survival rates for these patients range between 40-60%, but diagnosis after metastasis further reduces survival^5^. FLC tumors are notoriously refractory to standard chemotherapies and radiation treatments^6, 7^. Chemotherapy fails in most cases because tumor cells develop resistance to apoptosis, which leads to increased cancer invasion and progression to metastasis^8^. Thus, understanding the molecular mechanisms of chemotherapeutic resistance in FLC represents an important line of investigation in this burgeoning area of cancer research.

The DNAJ-PKAc fusion kinase is detected in >90% of FLC patients^3, 9^, with a few cases also bearing a PKA-RIα deletion^10^. Exome sequencing reveals a 400 kb deletion in chromosome 19 that results in translation of this fusion kinase, in which the chaperonin-binding domain of Hsp40 is fused in frame to exons 2-10 of PKAc^6, 9, 11^. Structural studies show that DNAJ-PKAc adopts the conformation of an active kinase with an operative J domain^12, 13^. While overwhelming evidence links this chimeric kinase to FLC, the molecular mechanisms that underlie DNAJ-PKAc action are unclear^3, 7, 14^. Identifying the pathological gains of function imparted by aberrant kinase activity or a new spectrum of protein-protein interactions that proceed through the DNAJ domain is important to understanding the signaling defects that occur in fibrolamellar carcinoma. More recently DNAJ-PKAc chimeras have been described in cholangiocarcinoma, hepatocellular carcinoma (HCC), and oncocytic biliary tumors^15–17^. Discovering druggable effectors and binding partners that function downstream of DNAJ-PKAc is a new strategy in the development of effective therapeutics for fibrolamellar carcinoma and other rare cancers.

Compartmentalization though association with A-kinase anchoring proteins (AKAPs) is a mechanism that imparts specificity to PKA signaling^18, 19^. AKAPs also integrate subcellular signals from other kinases and effector enzymes at sites proximal to their substrates^20–23^. Signaling through AKAPs limits the range and duration of information relay within cells^24–26^. In this report we show that DNAJ-PKAc interacts with a unique spectrum of binding partners^6, 14, 27^. Provocative new findings herein suggest that DNAJ-PKAc is largely excluded from AKAP signaling islands, thereby disrupting subcellular distribution of the enzyme. We find that substrate recognition of the chimeric kinase is unaffected. However, displacement from AKAP signaling islands affords the fusion enzyme access to aberrant substrates in cellular compartments not usually occupied by wildtype PKAc. One of these proteins is Bcl2-associated athanogene 2 (BAG2), a co-chaperone that is recruited to the fusion kinase through association with Hsp70. BAG2 has been linked to poor prognosis and resistance to chemotherapeutic agents in certain cancers^28–30^. Pharmacological studies with clinically relevant drugs implicate a pro-survival function of BAG2 at DNAJ-PKAc scaffolds and confirm a synergistic effect of etoposide and navitoclax on enhancing drug-induced cell death through Bcl-2 inhibition in cellular models of fibrolamellar carcinoma.

## RESULTS

### Enzyme-catalyzed proximity labeling identifies molecular associations with DNAJ-PKAc

Little is known regarding the molecular mechanism of DNAJ-PKAc in FLC pathogenesis. We first used enzyme-catalyzed proximity labeling to identify the interacting partners and protein components involved in aberrant DNAJ-PKAc complexes (Figure 1A). AML12 hepatocytes expressing PKAc variants tagged with the biotin ligase miniTurbo were generated using lentiviral transduction^19, 31, 32^. Single-cell clones were hand-picked and grown in individual wells of 48-well plates then screened for expression of the PKAc-miniTurbo variants using immunofluorescence staining (Figure 1B). Doxycycline-inducible expression of the miniTurbo fused bait proteins was adjusted to physiologically relevant levels of PKAc variants within the cells (Figure 1C, bottom). Neutravidin-HRP detection of biotinylated proteins demonstrates strong labeling for both cell lines after live incubation with biotin for 2 h (Figure 1C, top). Two clones each of DNAJ-PKAc- and WT PKAc-miniTurbo expressing cell lines were selected to proceed with proximity mass spectrometry. Quantitative analysis of the resulting LC/MS-MS data identified a total of 1174 proteins with 261 significant hits. Proteins with a log2(fold change) greater than 1.5 versus WT PKAc were labeled as either less associated (Figure 1D, red dots) or more associated (Figure 1D, blue dots) with the DNAJ-PKAc fusion. STRING analysis was used to create a network for proteins exhibiting greater association with DNAJ-PKAc. These include the co-chaperone proteins stress-induced phosphoprotein 1 (STIP1) and Bcl-2 associated athanogene 2 (BAG2; Figure 1E)^33^. Reactome pathway analysis of the DNAJ-PKAc proximitome using Enrichr revealed enrichment of pathways associated with mRNA processing and metabolism, as well as pathways associated with collagen biosynthesis and cytoskeletal remodeling that are involved in fibrosis and metastasis, respectively (Figure 1F)^34–39^.

**Figure 1.**
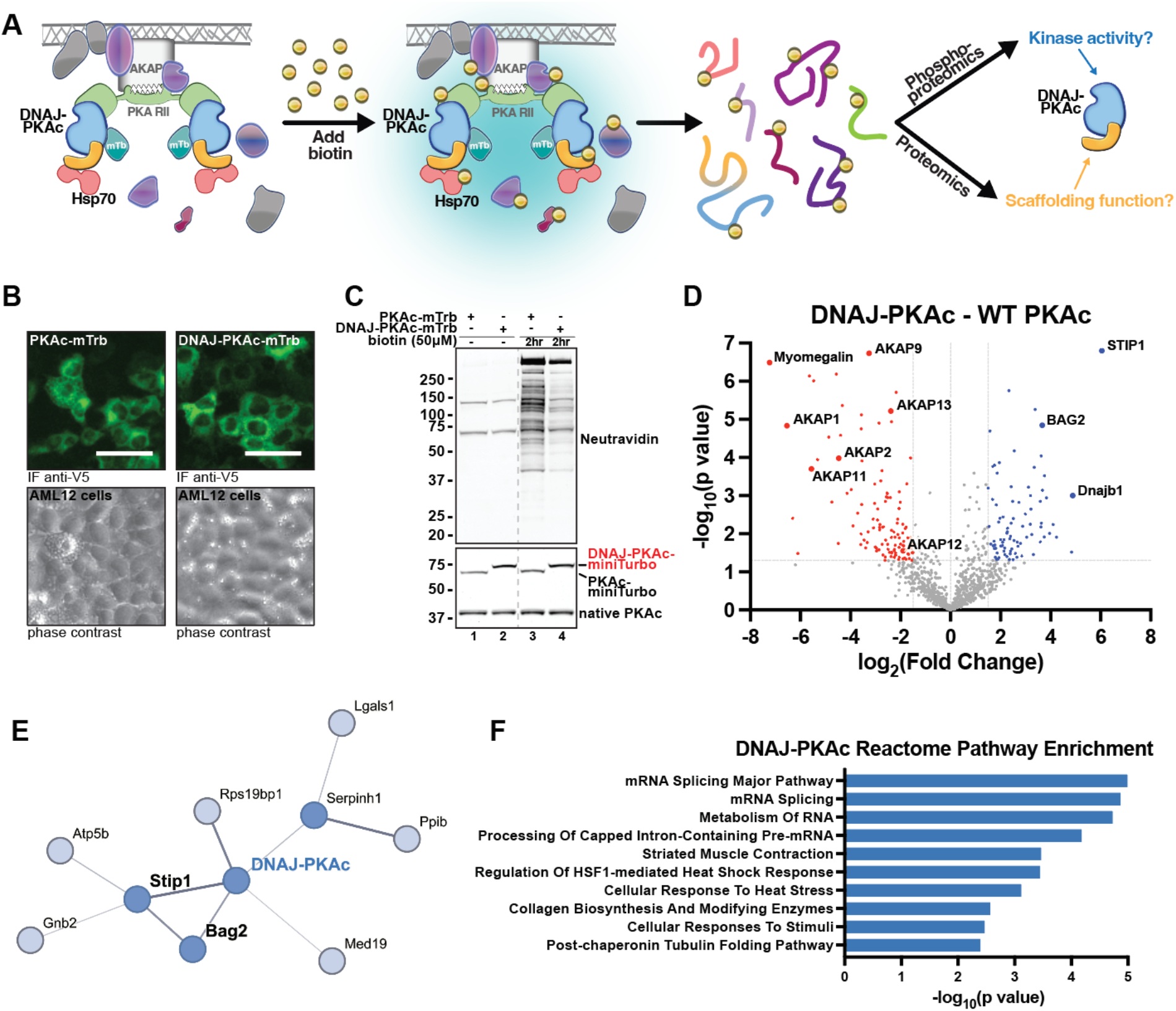
Enzyme-catalyzed proximity labeling identifies molecular associations with DNAJ-PKAc. A) Schematic of miniTurbo (mTrb) driven proximity labeling workflow and study design. Addition of biotin allows labeling of proteins within a 5-10 nm radius of bait protein. Following isolation, biotinylated or phosphorylated peptides were subject to mass spectrometry analysis. B) Immunofluorescence imaging of AML12 hepatocytes demonstrating inducible expression of PKAc-mTrb (green, top left) or DNAJ-PKAc-mTrb (green, top right) with corresponding phase contrast (bottom). Scale bar = 50 μm. C) Immunoblot of cell lysates from stable AML12 lines treated with either DMSO or biotin (50 μM). Neutravidin-HRP in top panel shows labeling of biotinylated proteins. PKAc in bottom panel shows expression of miniTurbo-tagged PKAc variants (top band) over native PKAc (bottom band). D) Volcano plot of mass spectrometry results showing proteins with increased (blue), decreased (red) association with DNAJ-PKAc compared to WT PKAc. Proteins with p value > 0.05 and/or log2(fold change) less than 1.5 are shown in grey. Six biological replicates. E) STRING network depiction of selected proteins with greater enrichment in DNAJ-PKAc versus WT PKAc. F) Bar chart of top ten enriched terms from the Reactome 2022 gene set library for proteins more associated with DNAJ-PKAc. Results are displayed based on -log10(p value).

### Mislocalization of DNAJ-PKAc from AKAP signaling islands

Our proximity proteomics screen revealed that AKAPs were generally less associated with the DNAJ-PKAc fusion. STRING analysis of proteins identified as having decreased association with DNAJ-PKAc revealed a network of AKAPs (Figure 2A). Quantitative analysis of peptide counts for both catalytic and regulatory subunits demonstrated reduced labeling of type II regulatory subunits by DNAJ-PKAc (Figure 2B). There was no significant difference in association with type I regulatory subunit alpha between WT PKAc and DNAJ-PKAc. Decreased peptide counts for AKAPs suggests displacement of DNAJ-PKAc from AKAP signaling islands (Figure 2C). To confirm this, photoactivation live-cell microscopy assays were performed to assess in situ localization of PKAc variants (Figure 2D). AML12 hepatocytes were transiently transfected with variants of PKAc fused to a photo-activatable (PA) mCherry (magenta), RIIα-iRFP (not shown), and AKAP79-YFP (cyan) to recruit PKAc holoenzymes to the plasma membrane. Following photoactivation of a 2 μm locus on the cell membrane, we observed that WT PKAc remains predominantly colocalized with membrane-associated AKAP79 (Figure 2D, top row; Figure 2E, gray trace; and Figure 2F, grey bar; Video S1). Conversely, DNAJ-PKAc rapidly diffuses into the cytosol, away from the membrane (Figure 2D, second row from top, Figure 2E, blue trace, and Figure 2F, blue bar; Video S2). Quantification of cells across three independent experiments demonstrates that DNAJ-PKAc exhibits increased mobility compared to WT PKAc (Figures 2E and 2F). In order to determine the mechanism of fusion kinase mislocalization, we repeated this assay with two additional controls, DNAJ^H33Q^-PKAc, a point mutation to disrupt Hsp70 binding, and PKAc^Δ14^, which mimics the N-terminal truncation of PKAc in the fusion protein^3, 40^. The PKAc^Δ14^ truncation remains localized at the plasma membrane via AKAP79 anchoring. This suggests the N-terminal portion of the kinase is not necessary for AKAP association (Figure 2D, bottom row, Figure 2E, green trace, and Figure 2F, green bar; Video S3). Interestingly, DNAJ^H33Q^- PKAc exhibits an intermediate phenotype, not demonstrating full retention at the plasma membrane but not diffusing completely into the cytosol (Figure 2D, second row from bottom, Figure 2E, pink trace, and Figure 2F pink bar; Video S4). Together, these experiments indicate that DNAJ-PKAc is not confined within AKAP signaling islands and that this effect is partially alleviated when Hsp70 association with the fusion kinase is disrupted.

**Figure 2.**
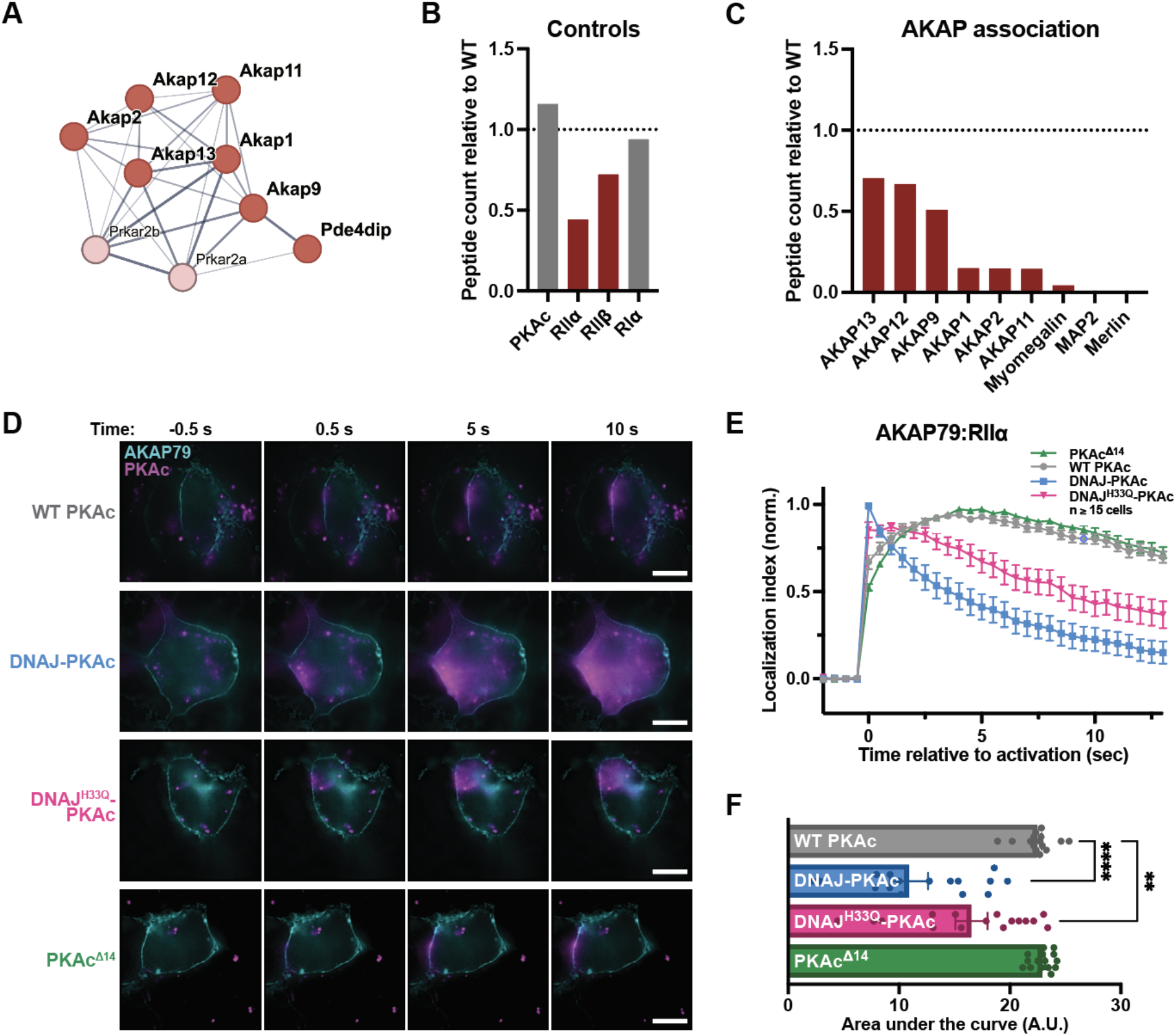
Mislocalization of DNAJ-PKAc from AKAP signaling islands. A) STRING network depiction of selected proteins with lesser enrichment in DNAJ-PKAc versus WT PKAc. B-C) Quantification of DNAJ-PKAc association with components of PKAc holoenzyme (B) and AKAPs (C) based on peptide count relative to WT PKAc. D) Photoactivation timecourses of AML12 hepatocytes expressing AKAP79-GFP, RIIα-iRFP, and either WT PKAc, DNAJ-PKAc, DNAJ^H33Q^-PKAc, or PKAc^Δ14^ tagged with photoactivatable mCherry. Scale bar = 10 μm. E) Quantification of (D). Data represents three experimental replicates. F) Integration of (E). Analyzed by one-way ANOVA and multiple comparisons corrected with Dunnett’s method; Mean ± SEM, ****p≤0.0001, **p≤0.01.

### Proximity phosphoproteomics uncovers distinct DNAJ-PKAc phosphorylation patterns

We next performed proximity phosphoproteomic enrichment screens to identify differences in substrate phosphorylation that occur due to mislocalization of DNAJ-PKAc. Following live biotin incubation and harvest, biotinylated peptides were isolated from stable AML12 cells expressing either WT PKAc-, DNAJ-PKAc-, or DNAJ-PKAc^K72H^-miniTurbo. Each sample was selectively enriched further for phosphorylated peptides (Figures 3A). Labeling efficiency and expression levels of each PKAc variant were verified prior to extraction and mass spectrometry analysis (Figures 3B, S3A, and S3B). Phospho-enriched mass spectrometry results comparing the fusion to WT PKAc identified 385 significant phosphopeptides of which 186 were less associated (Figure 3C, red dots) and 209 were more associated (Figure 3C, blue dots) with DNAJ-PKAc. These data showed trends in AKAP mislocalization (Figure S3C) and chaperone/co-chaperone association similar to those observed in the proximity proteomic screen (Figure 1D). Examination of the STRING network for phosphopeptides with increased DNAJ-PKAc association revealed gene ontology (GO) enrichment of biological processes often increased in cancer (i.e. mRNA processing, ribosome biogenesis, and translation), as well as members of the TORC1 signaling complex (Rps6, Akt, and Raptor; Figure S3D)^41–44^. Network propagation was performed on both the proximity proteomic (node fill color) and proximity phosphoproteomic (node border color) datasets to account for underlying network topology^45^. Integrated analysis of the two propagated proximity proteomic networks uncovered a cluster of significantly phosphorylated ribosomal proteins involved in translation, as well as a cluster of AKAPs that were both less associated and less phosphorylated in the presence of the oncogenic fusion kinase (Figure 3D).

**Figure 3.**
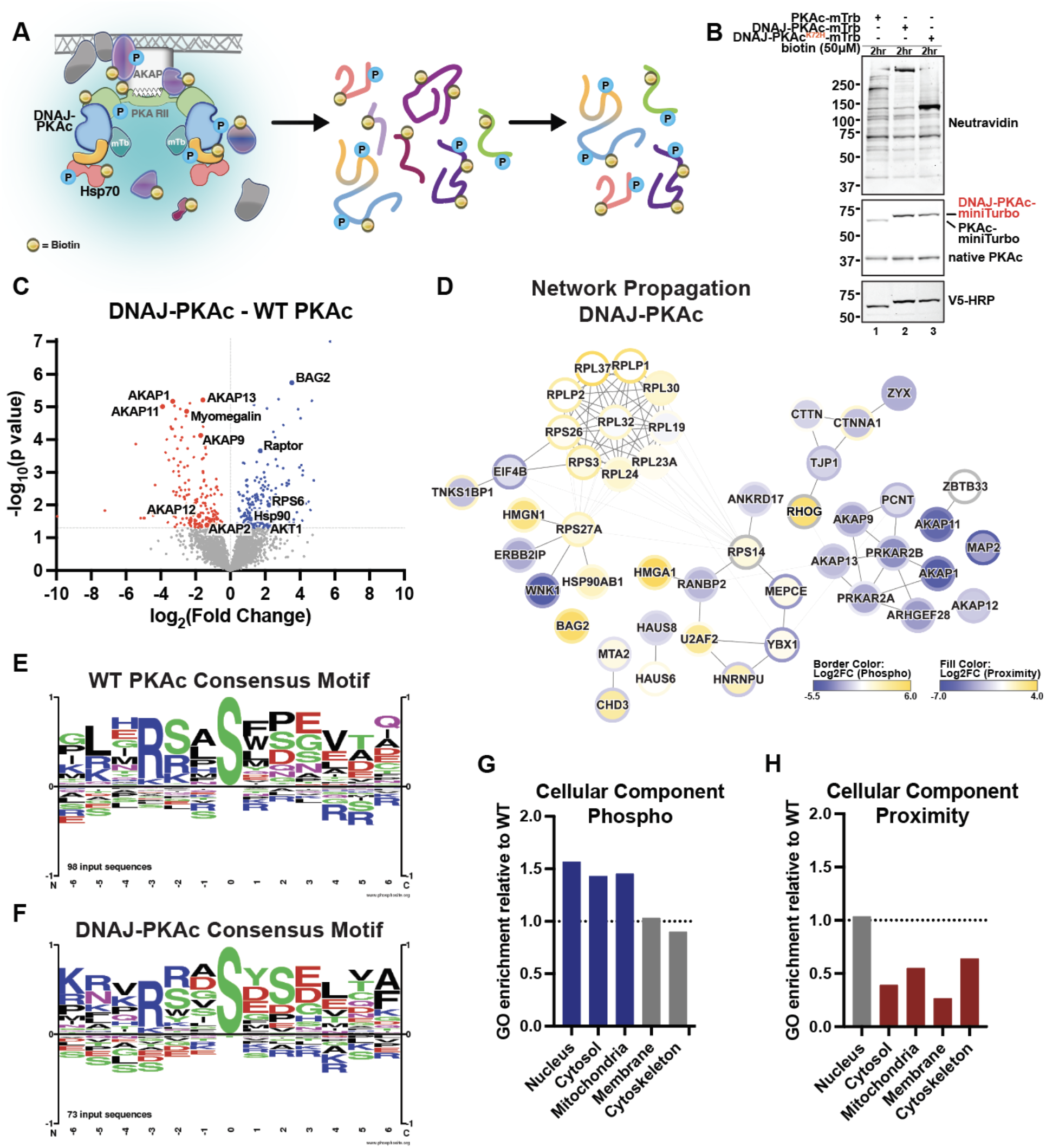
Proximity phosphoproteomics uncovers distinct DNAJ-PKAc phosphorylation patterns. A) Schematic of phosphoenrichment following miniTurbo-driven biotin labeling. B) Immunoblot of cell lysates from stable AML12 lines following biotin (50 μM) incubation. Neutravidin-HRP in top panel shows labeling of biotinylated proteins. PKAc in middle panel shows expression of mTrb-tagged PKAc variants (top band) over native PKAc (bottom band). V5-HRP in bottom panel shows specific expression of PKAc-mTrb variants. C) Volcano plot of mass spectrometry results showing phosphopeptides with increased (blue), decreased (red) association with DNAJ-PKAc compared to WT PKAc. Proteins with p value > 0.05 and/or log2(fold change) less than 1.5 are shown in grey. Four biological replicates. D) Depiction of results from ReactomeFI network propagation. Proximity proteomics is respresented by node fill color. Proximity phosphoproteomics is represented by node border color. Degree of association is represented on a yellow (more associated) to blue (less associated) spectrum based on log2(fold change). FDR < 0.1. See also Figure S3E. (E-F) Logo analysis depicting basophilic substrate consensus motifs for WT PKAc (E) and DNAJ-PKAc (F). (G-H) Gene ontology (GO) enrichment scores for DNAJ-PKAc cell components relative to WT PKAc.

Comparison of substrate motifs through alignment and logo analysis of basophilic phosphopeptides revealed only minor differences in kinase consensus sequences between WT PKAc and DNAJ-PKAc (Figure 3E and 3F). We next performed quantitative comparison of WT PKAc and DNAJ-PKAc GO enrichment scores for cellular component association. This revealed increased phosphorylation of proteins localized to the nucleus, the cytosol, and mitochondria in cells expressing DNAJ-PKAc (Figure 3G). Importantly, these phosphorylation events occurred despite the chimeric fusion having near equivalent association with proteins in the nucleus, and over 40% less association with proteins in the cytosol and mitochondria (Figure 3H). Thus, while substrate selectivity of the fusion kinase is not substantially altered, mislocalization from AKAP signaling complexes allows DNAJ-PKAc aberrant access to substrates in different cellular compartments than those occupied by WT PKAc.

### Substrates of DNAJ-PKAc are regulators of ribosome biogenesis and translation

To investigate changes in phosphorylation patterns due to DNAJ-PKAc, we compared the DNAJ-PKAc and kinase-dead DNAJ-PKAc^K72H^ proximity phosphoproteomic datasets. Analysis identified 536 significant phosphopeptides, 239 of which had increased association and 297 of which had decreased association with the DNAJ-PKAc active fusion compared to the kinase-dead fusion (Figure 4A). Examination of the phosphopeptides that were significantly more associated with DNAJ-PKAc when compared both to WT PKAc (Figure 3C, blue dots) and DNAJ-PKAc^K72H^ (Figure 4A, blue dots) uncovered 67 phosphorylation sites across 53 putative substrates of the FLC fusion kinase (Figure 4B). STRING network and GO term analysis of this phosphoprotein subset revealed increased association of DNAJ-PKAc with proteins involved in ribosome biogenesis and translation (Figure 4C). Peptides from regulators of these processes were at least two-fold more phosphorylated in the presence of the DNAJ-PKAc fusion versus WT PKAc (Figures 4D and 4E). A puromycin-based incorporation assay was performed to assess global translation activity in AML12 cells expressing DNAJ-PKAc (Figures 4F and S4). Compared to WT AML12 cells, AML12^DNAJ-PKAc^ cells showed significantly increased levels of protein synthesis (Figure 4G). This finding was further validated using a fluorescent reporter construct containing an IRES-linked GFP and mCherry. Results of this latter assay again demonstrated increased translational activity in cells expressing the DNAJ-PKAc fusion kinase (Figure 4H). Hence, the presence and activity of DNAJ-PKAc impacts processes involved in the translational capacity of the cell.

**Figure 4.**
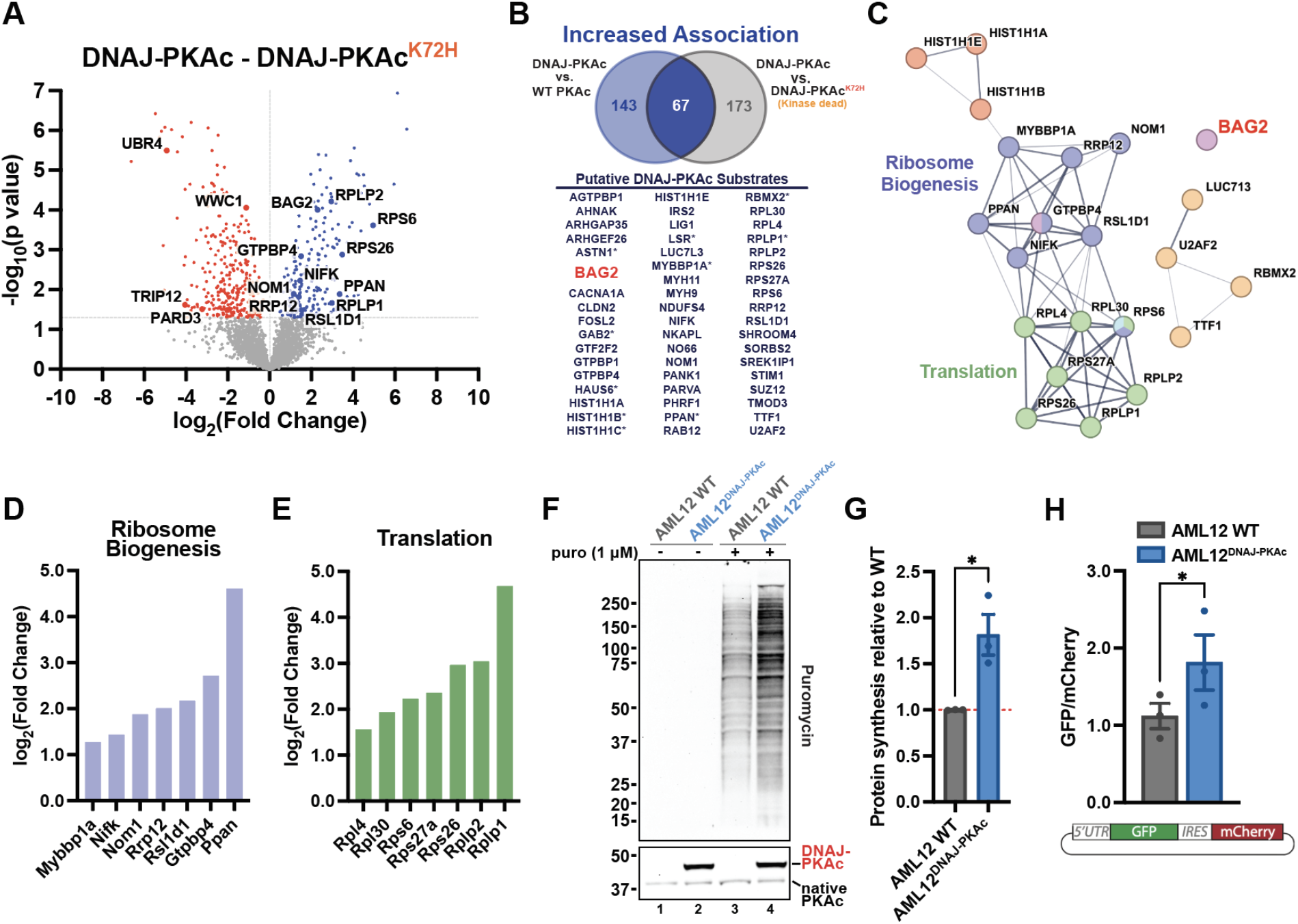
Substrates of DNAJ-PKAc are regulators of ribosome biogenesis and translation. A) Volcano plot of mass spectrometry results showing phosphopeptides with increased (blue), decreased (red) association with DNAJ-PKAc compared to DNAJ-PKAc^K72H^. Proteins with p value > 0.05 and/or log2(fold change) less than 1.5 are shown in grey. Four biological replicates. B) Venn diagram with resulting list of putative DNAJ-PKAc substrates identified by overlapping phosphosites that have increased association with DNAJ-PKAc versus WT PKAc and DNAJ- PKAc^K72H^. Asterisk indicates two or more phosphosites identified on corresponding protein. C) STRING network depicting selected functional clusters of putative DNAJ-PKAc substrates. D-E) Bar graphs showing log2(fold change) over WT PKAc for DNAJ-PKAc phosphoproteins associated with ribosome biogenesis (D) and translation (E). F) Immunoblot of cell lysates from WT AML12 and AML12^DNAJ-PKAc^ treated with either vehicle or puromycin (50 μM). Puromycin in top panel shows newly synthesized, puromycin-labeled proteins. PKAc in bottom panel shows expression of miniTurbo-tagged PKAc variants (top band) over native PKAc (bottom band). See also Figure S4. G) Quantification of (F) measuring protein synthesis in AML12^DNAJ-PKAc^ cells versus WT AML12 cells. Data represents three biological replicates. Mean ± SE. *p≤0.05. Fluorescent protein translation measurements (GFP intensity) in AML12 cells (grey) and AML12^DNAJ-PKAc^ cells (blue). Student’s t-test. Mean ± SE. *p≤.05.

### BAG2 is recruited to DNAJ-PKAc and overexpressed in FLC tumors

BAG family proteins are involved in a variety of important cellular functions including cell survival and stress response^28^. BAG2 is a known regulator of Hsp70-mediated protein refolding and CHIP-mediated ubiquitination that has been implicated in several different cancers^30, 46–49^. Our proximity-labeling data uncovered BAG2 as an interacting partner and putative substrate of DNAJ-PKAc (Figure 5A). Assessment of the BAG2 phosphopeptides identified through mass spectrometry revealed a basophilic kinase recognition motif surrounding the phosphorylated serine 20 (Figure 5B). The canonical PKA motif R-X-X-S/T* at this site is shared with MAPKAPK2, a p38 MAPK substrate which is known to phosphorylate BAG2 at Ser-20^50–52^.

**Figure 5.**
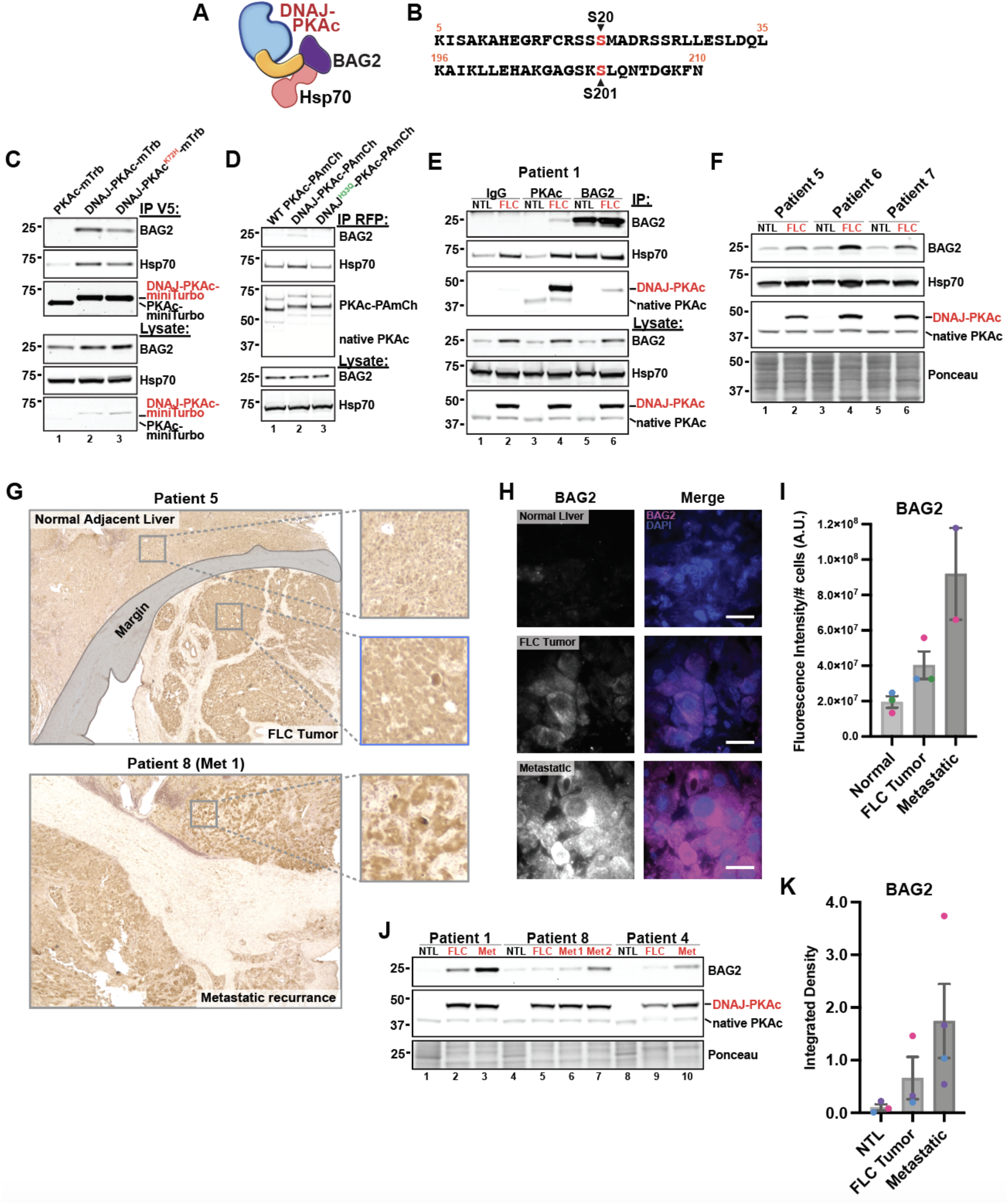
BAG2 is recruited to DNAJ-PKAc and overexpressed in FLC tumors. A) Model of BAG2 interaction with Hsp70 and DNAJ-PKAc. B) Peptide sequences of BAG2 phosphosites identified in MS screen. Phosphorylated residue is labeled in red. C) Immunoprecipitation of V5-tagged PKAc variants from stable AML12 cell lines. Represents three replicate experiments. D) Immunoprecipitation of mCherry-tagged PKAc variants from HEK293T cells. Represents three replicate experiments. E) Immunoprecipitation with antibodies to IgG control (lanes 1 and 2), PKAc (lanes 3 and 4), and BAG2 (lanes 5 and 6) from paired normal adjacent liver (NTL) and FLC tumor (FLC) tissue lysates from a single patient. F) Immunoblot of paired normal adjacent liver (NTL) and FLC tumor (FLC) tissue lysates from three patients. G) Chromogenic immmunohistochemistry of BAG2 in resected liver tissue from FLC patients. Images are representative of 8 normal liver/FLC tumor pairs and 2 metastatic tumors. H) Immunofluoresence staining of BAG2 in resected liver tissue from FLC patients. Scale bar = 20 μm. I) Quantification of (H). BAG2 fluorescence intensity normalized to number of cells in field. 10 images each per patient sample. Mean ± SE. J) Immunoblot of paired normal adjacent liver (NTL), FLC tumor (FLC), and metastatic tumor (Met) tissue lysates from three patients. K) Quantification of (J). Mean ± SE.

We next validated BAG2 recruitment to DNAJ-PKAc biochemically. Immunoprecipitation of PKAc variants from AML12 cells revealed that both active DNAJ-PKAc and the kinase-dead mutant co-precipitate BAG2 (Figure 5C, lanes 2 and 3) whereas WT PKAc does not (Figure 5C, lane 1). As it is known that BAG2 can associate with substrates both directly and indirectly via Hsp70, we wanted to determine if association of BAG2 was primarily Hsp70-mediated^46^. HEK293T cells were transiently transfected with either WT PKAc-, DNAJ-PKAc-, or DNAJ^H33Q^- PKAc-PAmCherry. Subsequent immunoprecipitation of each PKAc variant demonstrated that the DNAJ^H33Q^-PKAc mutant disrupted recruitment of not only Hsp70, but also BAG2, to the FLC fusion kinase (Figure 5D, lane 3). Importantly, immunoprecipitation of PKAc in 7 paired (normal liver and FLC primary tumor) and 4 metastatic patient tissue samples demonstrated that DNAJ-PKAc pulls down BAG2 in both primary (FLC) and metastatic (Met) tumors whereas WT PKAc in adjacent, non-tumor liver (NTL) does not (Figure 5E, lanes 3 and 4; Figures S5A and S5B). This result was further validated in a reciprocal immunoprecipitation of BAG2, which pulled down DNAJ-PKAc in FLC but not WT PKAc in neither FLC nor NTL tissue (Figure 5E, lanes 5 and 6).

BAG2 overexpression has been linked to poor clinical outcomes and prognosis in a variety of cancers^29, 30, 53, 54^. Therefore, we compared BAG2 levels in normal liver and primary tumor tissue within each patient. Immunoblot analysis of lysates from 7 different patients demonstrated overall increased expression of BAG2 in FLC versus NTL tissue (Figure 5F; Figures S5A, lysate panels). To visually assess BAG2 expression and global distribution in clinical samples, we performed immunohistochemistry on patient tissue sections. Chromogenic and immunofluorescent detection of BAG2 in stained tissue revealed a qualitative increase in BAG2 levels from normal adjacent liver to FLC tumor and primary tumor to metastasis (Figures 5G and 5F). Quantification of immunofluorescent images from 3 NTL/FLC pairs and 2 metastatic tissue sections confirmed this result (Figure 5I). Finally, evaluation of BAG2 expression via immunoblot in paired samples from 3 patients with advanced disease showed a progressive increase of BAG2 levels from non-tumor liver, to FLC primary tumor, to metastatic recurrence within each patient (Figures 5J and 5K). In sum, these findings suggest that BAG2 overexpression in FLC is correlated with disease progression.

### BAG2 promotes tumor cell survival and resistance to drug-induced cell death

BAG family proteins regulate cell growth, stress response, and cell death through interaction with several signaling partners, including Bcl-2, an inhibitor of cell death^55–57^. However, in tumorigenesis, the function of BAG2 with regard to cell death appears to be cancer-type-specific^54^. Therefore, we wanted to determine if BAG2 contributes to tumorigenesis and if its association with Bcl-2 protects against cell death in the context of FLC (Figure 6A). CRISPR/Cas9 gene editing was employed to generate a knockout (KO) of BAG2 in AML12 cells expressing the DNAJ-PKAc fusion (Figure 6B). It has been previously demonstrated that AML12^DNAJ-PKAc^ cells exhibit accelerated proliferation compared to WT AML12 cells^14^. Therefore, we assessed whether knocking out BAG2 in AML12^DNAJ-PKAc^ cells affects this fundamental property of tumorigenesis. Bromodeoxyuridine (BrdU) incorporation evaluated using a colorimetric ELISA showed reduced DNA synthesis in AML12^DNAJ-PKAc^ cells lacking BAG2 as compared to mock-control AML12^DNAJ-PKAc^ cells (Figure 6C). To establish the apoptotic role of BAG2 in FLC, we induced cell death in WT AML12, AML12^DNAJ-PKAc^, and AML12^DNAJ-PKAc^ BAG2 KO cells with the potent chemotherapeutic agent etoposide. Examination of cell density and morphology revealed that cells expressing DNAJ-PKAc were more resistant to etoposide-induced cell death as evidenced by minimal effects of treatment on cell growth, size, and shape compared to WT AML12 cells (Figure 6D, top panels). Conversely, BAG2 KO AML12^DNAJ-PKAc^ cells exhibited features such as swelling and blebbing, suggesting restored susceptibility to drug-induced cell death when subjected to etoposide (Figure 6D, bottom panels). A hallmark of cell death processes is PARP-1 activation and cleavage^58^. Immunoblot analysis and quantification of cleaved PARP-1 following etoposide treatment corroborated our findings, demonstrating higher levels of the 89-kDa apoptotic fragment in WT AML12 and AML12^DNAJ-PKAc^ BAG2 KO cells than in normal AML12^DNAJ-PKAc^ (Figures 6E and 6F).

**Figure 6.**
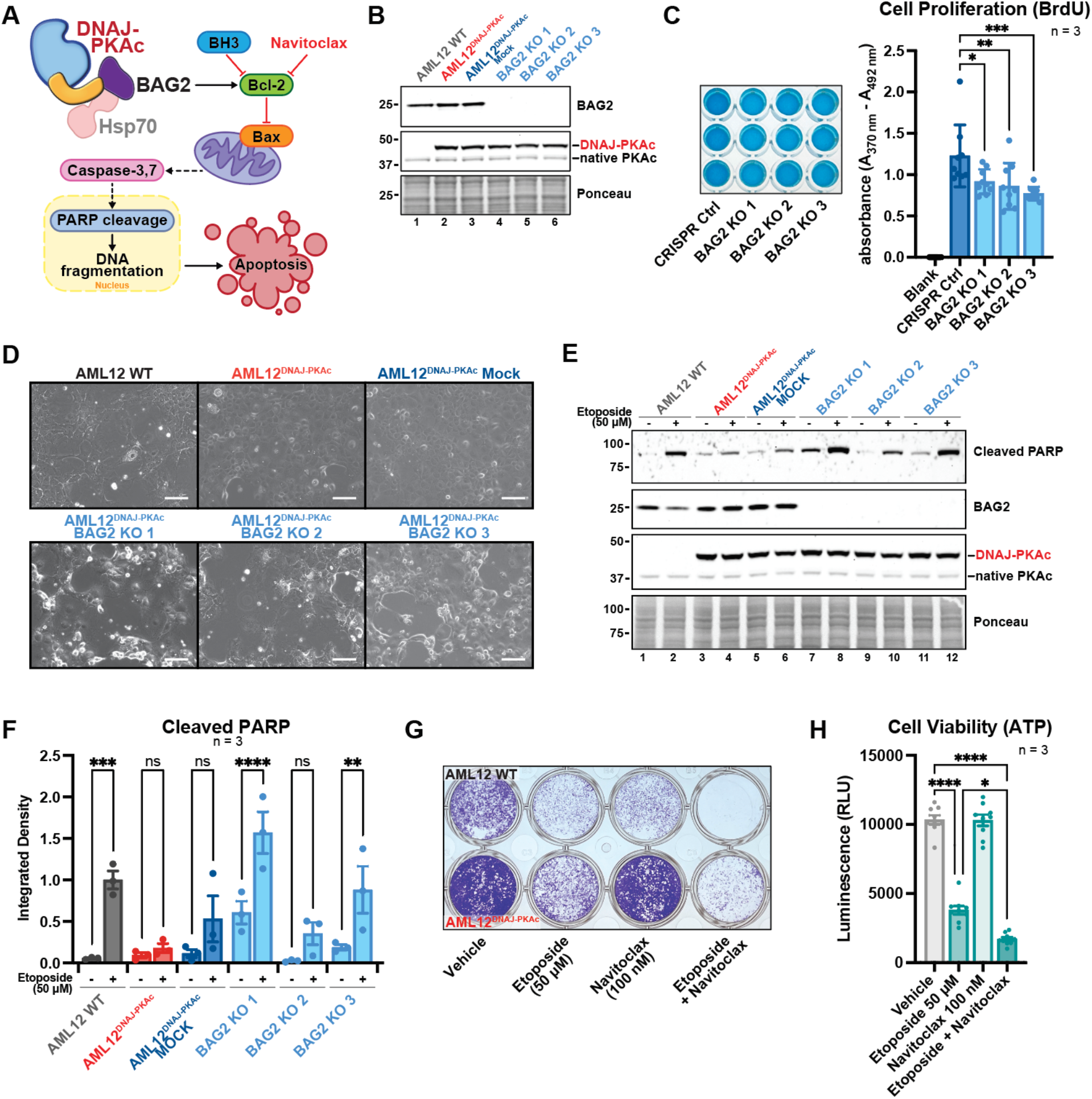
BAG2 promotes tumor cell survival and resistance to drug-induced cell death. A) Schematic showing BAG2 regulation of apoptosis in FLC. B) Immunoblot of three CRISPR/Cas9-generated BAG2 KO in AML12^DNAJ-PKAc^ clonal cell lines. C) ELISA BrdU incorporation assay measuring cell proliferation in three AML12^DNAJ-PKAc^ BAG2 KO clones versus AML12^DNAJ-PKAc^ Mock (CRISPR Ctrl) control line. Mean ± SE. ***p≤0.001, **p≤0.01, *p≤0.05. D) Phase contrast images showing etoposide-induced cell death in WT AML12, AML12^DNAJ-PKAc^, AML12^DNAJ-PKAc^ CRISPR Mock control, and three clonal AML12^DNAJ-PKAc^ BAG2 KO cell lines. E) Immunoblot of etoposide-induced PARP cleavage in WT AML12, AML12^DNAJ-PKAc^, AML12^DNAJ-PKAc^ CRISPR Mock control, and three clonal AML12^DNAJ-PKAc^ BAG2 KO cell lines. F) Quantitation of (E). Mean ± SE. ****p≤0.0001, ***p≤0.001, **p≤0.01. G) Crystal violet stain cell survival assay showing WT AML12 cells (top wells) and AML12^DNAJ-PKAc^ cells (bottom wells) treated with either vehicle, etoposide (50 μM), navitoclax (100 nM), or etoposide (50 μM) plus navitoclax (100 nM). Image is representative of 3 experimental replicates. H) Quantification of CellTiter-Glo luminescent assay showing cell viability of AML12^DNAJ-PKAc^ cells following treatment with either vehicle, etoposide (50 μM), navitoclax (100 nM), or etoposide (50 μM) plus navitoclax (100 nM). Data represents 3 experimental replicates. Mean ± SE. ****p≤0.0001, **p≤0.01.

Recently, BH3-mimetics have emerged as viable and potent targeted therapeutics in the context of cancer^59, 60^. These molecules, such as ABT-737/venetoclax and its derivative ABT- 263/navitoclax, inhibit Bcl-2 family proteins to induce apoptosis and are used clinically^61^. Therefore, we tested the effect of Bcl-2 inhibition on cell survival and susceptibility to induced cell death. Cytotoxicity was monitored by performing crystal violet staining after treatment with etoposide alone, navitoclax alone, or a combination of the two compounds (Figure 6F). WT AML12 cells were susceptible to both etoposide and navitoclax, and highly susceptible to the etoposide/navitoclax combination (Figure 6G, top wells), whereas cells expressing DNAJ-PKAc were moderately affected by etoposide treatment, completely resistant to navitoclax, and highly susceptible to the etoposide/navitoclax combination (Figure 6G, bottom wells). We further evaluated the impact of this drug combination using CellTiter-Glo to measure ATP levels, an indicator of cellular metabolism. Co-incubation with etoposide and navitoclax significantly and synergistically decreased AML12^DNAJ-PKAc^ cell viability more effectively than either compound alone (0.450 ± .042 versus etoposide, 0.163 ± .012 versus vehicle; SEM, n = 3) (Figure 6H). Hence, these studies implicate a pro-survival function of BAG2 in FLC, and pharmacologically confirm a synergistic effect of drug-induced cell death and Bcl-2 inhibition in this disease.

## DISCUSSION

Cancer is a genetic disease typically marked by mutations in 2-6 driver genes^62^. In contrast, the adolescent liver cancer fibrolamellar carcinoma emanates from a single genetic lesion on chromosome 19 that generates the chimeric enzyme DNAJ-PKAc^3, 63, 64^. Expression of a fusion kinase that encodes a chaperonin binding domain joined in frame with the catalytic core of PKAc is a dominant oncogenic event in FLC^3^. Yet, the molecular mechanisms by which this fusion kinase precipitates this aggressive and intractable liver cancer remain poorly understood. This is in part due to a scarcity of clinical samples available to investigate this rare tumor and limited access to animal models that faithfully recapitulate the molecular signature of FLC^64–67^. Here, we used live-cell enzyme-catalyzed biotinylation to identify DNAJ-PKAc-associated proteins and dysregulated biological processes in a hepatocyte model of FLC. While, aberrant kinase activity is assumed to be an initiating factor in this disease, the recruitment of DNAJ-PKAc specific binding partners, such as the chaperone Hsp70, implies that the scaffolding function of DNAJ-PKAc is another disease driving determinant^14, 68^. Abnormal enhancer activity, transcriptome remodeling, and altered translation have also been identified as factors contributing to FLC oncogenesis^27, 69^. Thus, a critical aspect of this work is the equivalent attention given to the catalytic actions, scaffolding functions, and downstream signaling impacts of this fusion kinase.

A-kinase anchoring proteins (AKAPs) are crucial for spatial regulation of protein kinase A and are responsible for directing the actions of this highly utilized multipurpose enzyme^18, 25, 70–73^. Surprisingly, our proximity mass spectrometry experiments identified a loss of DNAJ-PKAc association with AKAPs. This suggests FLC tumor cells harbor an unrestrained fusion kinase that increases off-target phosphorylation of cellular proteins. Interestingly, our live-cell photoactivation studies demonstrated that recruitment of the chaperone Hsp70 is a factor in the displacement of DNAJ-PKAc. Hsp70 often works as part of a chaperone/co-chaperone complex and therefore could mediate the association of other proteins with the catalytic subunit of PKA via the added J-domain^74^. Indeed, immunoprecipitation experiments in figure 5D demonstrate that the Hsp70- binding mutant of DNAJ-PKAc fails to bind the co-chaperone BAG2. Together, these data suggest that disruption of AKAP interaction could be a consequence of recruiting binding partners that impede the geometric organization required for PKAc anchoring. This unexpected result stands in contrast to a biochemical report finding no differences in the binding of PKAc and DNAJ-PKAc to regulatory subunits^68^. Thus, mislocalization of DNAJ-PKAc may be context dependent and subject to the availability of binding partners such as Hsp70 that block association with regulatory subunits or AKAPs.

Displacement of DNAJ-PKAc from AKAP complexes is reminiscent of Cushing’s adenomas, where distinct mutations in the catalytic core of PKAc disrupt AKAP-dependent compartmentalization^75^. In adrenal Cushing’s syndrome, mislocalization arises from perturbations to the protein-protein interface by which PKAc associates with regulatory subunits. In contrast the acquired scaffolding function of DNAJ-PKAc confers enhanced cytoplasmic mobility in FLC. Furthermore, precision medicine approaches have shown that adrenal Cushing’s stems from a range of mutations in different regions of the catalytic subunit whereas FLC normally arises from a single genetic lesion^68, 76^. Mutations in the catalytic core of Cushing’s kinases have been suggested to cause substrate rewiring that alters how the kinase recognizes basophilic substrate motifs^76–78^. In contrast, our analyses of proximal phosphorylated peptides in figure 3E and F indicate that native PKAc and the fusion kinase utilize virtually identical recognition motifs. Thus, we argue that displacement from AKAPs results in altered compartmentalization of DNAJ-PKAc, providing this promiscuous kinase access to additional compatible substrates. For instance, increased phosphopeptides from TORC1 complex components AKT1 and Raptor echo previous reports of mTOR involvement in FLC, where hyperphosphorylation of S6K was observed^79^.

Moreover, three of the largest biological process clusters consisted of mRNA processing, ribosome biogenesis, and protein translation machinery (Figure S3D). This finding is consistent with the involvement of DNAJ-PKAc signaling in translation initiation, an oncogenic process effectively targeted using a clinically-available eif4a inhibitor in a patient-derived cellular model of FLC^27^.

A troubling feature of fibrolamellar carcinoma is its resistance to systemic chemotherapies^80^. Our study presents a likely mechanism for this challenging complication through the recruitment of BAG2. This co-chaperone belongs to a family of proteins whose key functions center around the regulation of cell survival, apoptosis, and stress response^81, 82^. Several studies have investigated the oncogenic action of BAG2 and proposed context specific roles for this co-chaperone. A study in thyroid cancer found that BAG2 adopts a pro-apoptotic function following proteasome inhibition^83^. Conversely, recent studies have identified a more prevalent role for BAG2 in promoting cell survival and tumorigenesis. For example, high levels of BAG2 in gastric cancer are linked to poor outcomes^84^. Furthermore, BAG2 upregulation in both glioma and breast cancer confers chemoresistance and protection against apoptosis^29, 49, 53, 85^. Finally, research in hepatocellular carcinoma reports decreased overall survival in patients with elevated BAG2^54, 86^. Remarkably, each study describes a different strategy by which BAG2 contributes to disease progression. However, a common thread, regardless of mechanism, is that increased BAG2 expression is correlated with poor disease prognosis. In FLC patient tissue, we find that BAG2 protein levels are elevated with severity of disease, increasing first in liver tumors, and peaking in metastases of the advanced cancer. In this way, BAG2 may be considered a biomarker for disease progression in FLC.

As outlined above, the molecular mechanism of BAG2 action in tumorigenesis is multi- faceted. Emerging data has linked BAG2 to elements of MAPK signaling in the context of cancer^84, 86^. BAG2 is directly phosphorylated at serine 20 by MAPKAPK2, a major downstream mediator of p38-dependent processes^52^. Alterations in p38/MAPK pathway have been shown to contribute to tumor growth and metastatic progression in certain cancers^87^. In addition, association with Bcl-2, an inhibitor of the pro-apoptotic protein Bax, suggests an anti-apoptotic role for BAG family proteins^88, 89^. Indeed, recent studies have established BAG2 as a promoter of cell survival in the regulation of apoptosis^84, 90, 91^. Our results demonstrate that BAG2 is not only more associated with the fusion kinase, but also more phosphorylated (at serine 20) in the presence of DNAJ-PKAc (Figures 3D and 5B). We further propose that upregulation of BAG2 may contribute to chemotherapy resistance through attenuation of cellular apoptosis. Enhanced sensitivity to etoposide-induced cell death in FLC model cells lacking BAG2 supports this mechanism (Figure 6D and 6E). This also led to our hypothesis that intervention at the level of Bcl-2 with the BH3-mimetic navitoclax may circumvent BAG2-mediated suppression of and resistance to apoptosis. Both compounds have been clinically evaluated. Etoposide is an FDA-approved topoisomerase II inhibitor used alone or in combination for the treatment of various cancers including small cell lung cancer, testicular cancer, and lymphoma^92^. Navitoclax, the predecessor to the FDA-approved Bcl-2 inhibitor venetoclax, is currently in Phase III clinical trials following promising Phase II results as a combination therapy in myelofibrosis^93, 94^. The success of the combination drug experiments in our model cell line offers promise for therapeutically targeting apoptotic-resistance in FLC tumors.

In conclusion, we have discovered that the pathogenic PKA fusion kinase driving fibrolamellar carcinoma is displaced from AKAP signaling islands, leading to abnormal phosphorylation of substrates residing in distal subcellular compartments. Additionally, we have established that DNAJ-PKAc drives enhanced cell survival through interaction with BAG2, and that this oncogenic mechanism can be sensitized to pro-apoptotic drugs via BAG2 deletion or inhibition of Bcl-2. Future studies will undoubtably focus on discerning the therapeutic value of targeting the DNAJ-PKAc/BAG2/Bcl-2 axis in fibrolamellar carcinoma.

## ACKNOWLEDGMENTS

The authors would like to thank K. Collins, K. Jones, and J. Nelson for technical assistance and helpful discussions. This work was supported by NIH grants T32GM775043 (S.M.L), DK119186 (J.D.S.), and DK119192 (J.D.S.), a research grant from the Fibrolamellar Foundation (J.D.S.). The authors declare no conflicts of interest.

## MATERIALS AND METHODS

### EXPERIMENTAL MODEL AND SUBJECT DETAILS

#### Human liver tissue

Human FLC tumor and paired normal liver tissue were obtained in collaboration with the Yeung Lab in the UW Department of Surgery. De-identified patient tissue samples were either formalin-fixed, paraffin-embedded and mounted on glass slides for imaging, or fresh-frozen and stored at -80°C until homogenization for either immunoprecipitation in lysis buffer containing 2 mM EDTA, 20 mM NaF, 130 mM NaCl, 50 mM Tris pH 7.5 (at 4°C), and 1% Triton X-100, with protease inhibitors (10 μM leupeptin/pepstatin, 1 mM benzamidine, and 1 mM AEBSF) and phosphatase inhibitor (sodium β-glycerophosphate), or western blot in RIPA lysis buffer containing (1% NP-40 Tergitol, 0.5% deoxycholate, 0.1% SDS, 130 mM NaCl, 20 mM NaF, 2 mM EDTA, and 20 mM Tris pH 7.5 (at 4°C) along with 1 mM AEBSF, 10 μM leupeptin/pepstatin, and 1 mM benzamidine).

#### Cell lines and cell culture

HEK293T cells for lentiviral production were obtained from GE Lifesciences and maintained in DMEM containing 10% Gemini FBS. Wildtype AML12 hepatocytes were obtained from the Riehle Lab via ATCC and were developed by the Nelson Fausto lab (Wu et al., 1994). AML12^DNAJ-PKAc^ cells were generated previously by Rigney Turnham (Turnham et al., 2019). All AML12 cell lines were maintained in DMEM/F12 with 10% Gemini FBS, 1 mL 500X ITS supplement (Lonza 17- 838Z; 5 μg/mL insulin, 5 μg/mL transferrin, 5 ng/mL selenium), 50 μg/mL gentamycin, and 0.1 μM dexamethasone. All cell lines used in this study were grown at 37°C with 5% CO2.

#### Microbe strains

Amplification of non-viral mammalian expression plasmids was performed in GC10 competent cells (Genesee) and grown at 37°C. Amplification of viral vectors was performed in either Stbl3 (Invitrogen) or Stable (NEB) competent cells and grown at 30°C.

### METHOD DETAILS

#### Antibodies

The following antibodies were used in our studies: V5-tag Thermo Fisher R96025 (IF, IP); PKAc BD 610981 (IF, WB, IP); NeutrAvidin-HRP Pierce 31030 (WB); Puromycin Millipore (MABE343); PKAc CST 5842 (WB); BAG2 Invitrogen PA5-78853 (IF, WB, IP, IHC); Hsp70 Proteintech 10995- 1-AP (WB); RFP Rockland 200-101-379 (IP); RFP GenScript A00682 (WB); Cleaved PARP CST 94885 (WB).

#### Plasmid generation

Specific plasmids are listed in the key resources table. Standard cloning was performed using PCR (35 μL ddH2O, 10 μL 5x HF Phusion buffer (NEB), 1 μL of 10 mM mixed dNTPs, 2.5 μL combined primers at 10 mM each, 1 μL template DNA at 10 ng/mL, and 0.5 μL Hot-Start Phusion polymerase (NEB)) in a Bio-Rad thermocycler. Thermocycling protocols varied depending on primer conditions and length of target region (30 s/kb). For mutagenesis protocols, DpnI restriction enzyme and polynucleotide kinase were used (NEB). Some constructs were made using the Gateway cloning system (Thermo Fisher). Ligation was performed with T4 DNA ligase (NEB) for 10-20 min at RT or at 4°C overnight using manufacturer’s recommendations. Transformation into competent DNA (see *Microbe strains* above) was performed on ice for 15-30 min before heat shock for 30 s at 42°C.

#### Generation of BAG2 KO AML12^DNAJ-PKAc^ cells using CRISPR/Cas9 gene editing

A pool of vectors encoding 3 different Bag2-specific gRNAs (sequences available from manufacturer upon request), Cas9 enzyme, and GFP were purchased from Santa Cruz Biotechnology, Inc (Dallas, TX). AML12 hepatocytes containing the DNAJ-PKAc mutation were transfected with the pooled plasmids using Lipofectamine 3000 Reagent. 24 h after transfection, single cells were sorted according to GFP fluorescence using FACS into 96-well plates. Clones were screened by immunoblot for loss of BAG2 expression.

#### Immunofluorescent staining

Cells with inducible expression of miniTurbo-fused PKAc subunit variants were plated in 48 well tissue culture dishes. 16-24 h later, doxycycline was added to induce overexpression of bait proteins. 48 h post-induction, cells were fixed with 4% paraformaldehyde in PBS for 15 min at 25°C and washed 3x with PBS. Cells were then blocked for 1 h at 25°C in 3% BSA and 0.3% Triton X-100 in PBS and primary antibodies diluted in blocking solution were applied to the cells at 4°C for 12-16 h. Following 3x washes with PBS, cells were incubated at 25°C with fluorescent secondary antibodies (used at 1:1000) and DAPI (∼1:10,000). Cells were washed three more times with PBS and imaged on a Keyence BZ-X710 microscope.

#### Immunoblotting

Cell lysates were made using RIPA lysis buffer (1% NP-40 Tergitol, 0.5% deoxycholate, 0.1% SDS, 130 mM NaCl, 20 mM NaF, 2 mM EDTA, and 20 mM Tris pH 7.5 (at 4°C) along with 1 mM AEBSF, 10 μM leupeptin/pepstatin, and 1 mM benzamidine). Ηuman liver protein extracts were made by homogenizing fresh frozen tissue sections in RIPA buffer. For experiments to detect S/T phosphoproteins, 10 mM β-glycerophosphate was added. Samples were incubated 5 min on ice and spun at 15,000 x *g* for 10 min at 4°C. Protein concentration was measured by BCA (Thermo Scientific). Gels were loaded with 15-20 mg protein after heating for 10 min at 80°C with PAGE sample buffer containing 3% (final) β-mercaptoethanol. Proteins were transferred to nitrocellulose, incubated with ponceau S to measure total protein loading, blocked in 5% milk TBST for at least 30 min at RT, and probed with antibodies in 5% BSA TBST or 5% milk TBST overnight at 4°C. Membranes were washed 3 times in TBST and then incubated with secondary antibodies conjugated to HRP diluted in 5% milk TBST for 1-2 h at RT. Following secondary antibody incubation, membranes were washed again 3 times in TBST and signals were visualized with SuperSignal West Pico Chemiluminescent Substrate (Thermo Fisher) on an Invitrogen iBright FL1000 Imaging System. Quantification was performed with ImageJ analysis software (FIJI) by measuring signal minus background for each band and dividing by the appropriate control signal, as indicated in each figure.

#### Proximity biotinylation and sample prep for MS

Stable AML12 cell lines were made using lentivirus encoding a tetracycline-responsive promoter and variants of PKAc tagged with V5 and miniTurbo biotin ligase at the C-terminus. Doxycycline (0.5-1 μg/mL) was used to induce optimized overexpression of the bait constructs, as determined by PKAc immunoblotting. miniTurbo-tagged variant expression was induced for 48 h prior to application of 50 μM biotin in DMSO. Cells were incubated for 2 h at 37°C, washed 2 times for 1 min using 10 mL PBS to deplete excess biotin, and then lysed using RIPA buffer (as described above). Protein concentrations were measured by BCA and samples were diluted to 1 mL of 0.5 mg/mL in RIPA buffer in low protein binding collection tubes (Thermo Fisher) containing 25 μL of NanoLink® magnetic streptavidin beads. Tubes were rotated 1 h at RT and placed on a magnet. Supernatant was saved for diagnostics and samples were washed in RIPA 2 times, 2 M urea in 20 mM Tris 2 times, and 25 mM Tris 2 times. For normal mass spectrometry analysis, samples were resuspended in 8 M urea in 100 mM Tris pH 8.5 with 5 mM tris(2-carboxyethyl)phosphine hydrochloride (TCEP) and 10 mM chloroacetamide (CAM) and then incubated at 37°C for 1 h. For phosphopeptide mass spectrometry analysis, samples were resuspended in 20% trifluoroethanol 25 mM Tris pH 7.8 with 5 mM TCEP and 10 mM CAM and incubated at 95°C for 5 min. For digestion, samples were diluted 2-fold with 100 mM TEAB and 1 μg LysC was added before incubation for 2 h shaking at 37°C. Samples were again diluted with 100 mM TEAB and 1 μg Trypsin was added before incubation overnight shaking at 37°C. In the morning, normal mass spectrometry samples were acidified to 1% formic acid and loaded on C18 StageTips. Samples for phosphoproteomics were subjected to phosphopeptide enrichment using a Thermo Scientific High-Select™ Fe-NTA Phosphopeptide Enrichment Kit prior to StageTip loading.

#### LC-MS analysis

Peptides were eluted from StageTips using elution buffer (40% acetonitrile, 1% FA) and then loaded on a self-pulled 360 mm OD x 100 mm ID 20 cm column with a 7 mm tip packed with 3 mm Reprosil C18 resin (Dr. Maisch, Germany). For pull-down experiment, peptides were analyzed by nanoLC-MS in a 90 min gradient from 15% to 38% buffer B (for phosphopeptides 6%–35% buffer B) at 300 nL/min using a Thermo EASY nLC 1200 system (buffer A: 0.1% acetic acid; buffer B: 0.1% acetic acid, 80% acetonitrile). Mass spectra were collected from an Orbitrap Fusion Lumos Tribrid Mass Spectrometer using the following settings. For MS1, Orbitrap FTMS (R = 60, 000 at 200 m/z; m/z 350–1600; 7e5 target; max 20 ms ion injection time); For MS2, Top Speed data-depen- dent acquisition with 3 s cycle time was used, HCD MS2 spectra were collected using the Orbitrap mass analyzer(R = 30,000 at 200 m/z; 31% CE; 5e4 target; max 100 ms injection time) an intensity filter was set at 2.5e4 and dynamic exclusion for 45 s.

#### Mass spectrometry data analysis

Mass spectra were searched against the UniProt human reference proteome downloaded on July 06th, 2016 using MaxQuant v1.6.2.6. Detailed MaxQuant settings: for phosphopeptide analysis, samples were set to fraction 1 and 5 for WT and mutant, respectively, to allow within-group ‘‘match between run’’; for pull-down, ‘‘Label-free quantification’’ was turned on, but not ‘‘match between run’’, no fractionation was set; Trypsin/P was selected in digestion setting. Other settings were kept as default. Protein network prediction and gene ontology analysis were performed using STRING database version 11.5 and gene ontology enrichment analysis was performed using The Gene Ontology Resource powered by PANTHER. Reactome pathway analysis was performed using Enrichr. For substrate motif predictions, PhosphoSitePlus® sequence logo analysis was performed on significantly enriched phosphosites for each PKAc variant.

#### Photoactivation assay

Wildtype AML12 hepatocytes were grown in glass bottom 35 mm dishes and transfected using Lipofectamine 3000 48 h before imaging. Mammalian expression plasmids with CMV promoters and encoding AKAP79-YFP, RIIα-iRFP, and either WT PKAc, DNAJ-PKAc, DNAJ^H33Q^-PKAc, or PKAc^Δ14^ tagged with photoactivatable mCherry were used. Imaging was performed using a GE OMX SR system. Exposure and laser intensity were optimized for each experimental replicate and held constant among experimental conditions. Photoactivation laser duration was kept under 50 milliseconds to activate a discrete area with minimal spread in the first image collected after activation. Images were collected at 2 Hz in 3 channels. A baseline of 4 images was taken prior to activation of the PKAc fluorophore. Cells were selected for imaging only when RIIα-iRFP signal was colocalized with AKAP79 signal. Secondary screening for this was performed posthoc. Timecourses were measured using ImageJ analysis software (FIJI). A localization index (intensity of the activated region divided by intensity of cytosolic region 6-8 mm distal) was used to interrogate change in fluorescent signal localization over time (mobility). For representative images and videos, deconvolution and alignment of green and far-red channels were performed using OMX software.

#### Puromycin translation assay

Cells were seeded at 400,000 cells/well in a 6-well plate. 48 h after plating, media was replaced with fresh media containing puromycin (1 μΜ) and cells were returned to 37°C incubator. After 30 min, cells were washed twice with DPBS for 1 min and cells were lysed in RIPA buffer (as described above) for immunoblot analysis.

#### Immunoprecipitation

Cell lysates were made using lysis buffer containing 1% Triton X-100, 130 mM NaCl, 20 mM NaF, 2 mM EDTA, and 50 mM Tris pH 7.5 (at 4°C) along with 1 mM AEBSF, 10 μM leupeptin/pepstatin, and 1 mM benzamidine. Lysates were incubated 5 min on ice and spun at 15,000 x *g* for 10 min at 4°C. Protein concentration was measured by BCA (Thermo Scientific) and adjusted to 0.5 mg/mL (or 1 mg/mL for human tissue using lysis buffer. Samples (500 mL) were precleared by rotating with 20 μL protein G agarose beads for 30 min at 4°C. Supernatants were then incubated with 1-2 μg of the appropriate antibody overnight. In the morning, 30 μL of protein G agarose beads were added and samples were returned to 4°C rotation for 1 h. Beads were washed with lysis buffer 3 times and centrifuged at 5000 x *g* following each wash, then aspirated with a 27G needle before resuspending in 1x PAGE sample buffer (3% β-mercaptoethanol, final) and heating at 80°C for 10 min. Figures are representative for at least 3 experimental replicates.

#### Immunohistochemistry

Formalin-fixed, paraffin-embedded normal, FLC, or metastatic liver tissue sections were deparaffinized by placing slides in 100% xylenes once for 10 min and once for 5 min. Samples were then rehydrated by placing slides in 100% ethanol twice for 10 min each, followed by 95% ethanol for 10 min, 80% ethanol for 10 min, and deionized water two times for 5 min each. Antigen retrieval was performed by placing slides in a chamber with pre-boiled 10 mM sodium citrate buffer (pH 6.0). The chamber was then placed inside of a vegetable steamer for 1 h. Slides were placed under cold running water for 10 min before permeabilization in 0.4% Triton X-100/PBS for 7 min. Blocking was carried out in 5% BSA and 10% donkey serum in PBST (containing 0.05% Tween) for 2 h at RT. <u>For fluorescent IHC</u>, slides were incubated with primary antibodies in 5% BSA in PBST overnight at 4°C. Cells were washed 3x in PBST for 10 min each and incubated with Alexa Fluor conjugated secondary antibodies and DAPI in 3% BSA in PBST for 1 h at RT. Slides were then washed six times for 10 min each in PBST. Samples were mounted on glass coverslips using ProLong® Diamond anti-fade mountant (Thermo Fisher) and cured overnight. Images were acquired using a GE OMX SR system. Signal intensity was measured using ImageJ analysis software (FIJI) and normalized to number of cells in image field (DAPI).

#### BrdU ELISA assay

Cells were seeded at 10,000 cells/well in a 96-well plate. Each condition was run in triplicate. Assay was optimized for cell type used and otherwise performed according to manufacturer’s protocol. 48 h after plating, BrdU labeling solution was added at 10 μΜ to each well and cells were returned to incubator. After 4 h, media in each well was replaced with FixDenat solution at 25°C for 30 min. FixDenat solution was then thoroughly removed and replaced by Anti-BrdU POD-conjugated antibody solution for 90 min at 25°C, followed by 3x wash with 200 μL PBS. 100 μL TMB substrate solution was then added to each well and color was allowed to develop over 30 min. Absorbance was read at 370 nm with a reference wavelength of 492 nm.

#### Etoposide-induced apoptosis assay

Cells were seeded at 200,000 cells/well in a 12-well plate. 48 h after plating, media was replaced with fresh media containing either DMSO or etoposide (50 μM) and cells were returned to 37°C incubator. 72 h later, cells were imaged on a Keyence BZ-X710 microscope, then washed once with DPBS and cells were lysed in RIPA buffer (as described above) for immunoblot analysis.

#### Cell viability assays

For crystal violet staining, cells were seeded at 50,000 cells/well in a 24-well plate. 48h after plating, media was replaced with fresh media containing either DMSO, etoposide (50 μM), navitoclax (100 nM), or a combination of the two and cells were returned to 37°C incubator. After 72 h, media was removed and replaced with crystal violet solution (0.25% crystal violet powder and 10% methanol in water) for 20 min. Cells were then washed 3 times with water and plate was allowed to dry for 24 h before imaging. For CellTiter-Glo, cells were seeded at 3,000 cells/well in a 96-well plate. Cells were allowed to recover for 16-24 h, then were treated either DMSO, etoposide (50 μM), navitoclax (100 nM), or a combination of the two and returned to 37°C incubator. After 72 h, CellTiter-Glo reagent was added and plate was placed on a dual-orbital shaker for 2 min to induce cell lysis. Plate was incubated at room temperature for 10 min and luminescence was recorded using a POLARstar Omega microplate reader.

### QUANTIFICATION AND STATISTICAL ANALYSES

Data quantification and statistical analyses as indicated in each figure legend were performed with GraphPad Prism 9 for Mac. All data are presented with mean ± SEM unless otherwise noted in the figure legend. Individual figure legends contain specific information on statistical parameters. Experiments involving more than three conditions used one-way ANOVA with subsequent t-tests corrected for multiple comparisons. Specific statistical approaches were determined based on the distributions and parameters for each dataset.

## Notes

### Competing Interest Statement

The authors have declared no competing interest.

## REFERENCES

1. Gallicchio, L., Daee, D.L., Rotunno, M., Barajas, R., Fagan, S., Carrick, D.M., Divi, R.L., Filipski, K.K., Freedman, A.N., Gillanders, E.M., et al. (2021). Epidemiologic Research of Rare Cancers: Trends, Resources, and Challenges. Cancer Epidemiol Biomarkers Prev 30, 1305–1311. 10.1158/1055-9965.EPI-20-1796.

2. Stransky, N., Cerami, E., Schalm, S., Kim, J.L., and Lengauer, C. (2014). The landscape of kinase fusions in cancer. Nat Commun 5, 4846. 10.1038/ncomms5846.

3. Honeyman, J.N., Simon, E.P., Robine, N., Chiaroni-Clarke, R., Darcy, D.G., Lim, II, Gleason, C.E., Murphy, J.M., Rosenberg, B.R., Teegan, L., et al. (2014). Detection of a recurrent DNAJB1-PRKACA chimeric transcript in fibrolamellar hepatocellular carcinoma. Science 343, 1010–1014. 10.1126/science.1249484.

4. Zack T, L.K., Maisel SM, Wild J, Yaqubie A, Herman M, Knox JJ, Mayer RJ, Venook AP, Butte A, O’Neill AF, Abou-Alfa GK, Gordan JD (2023). Defining incidence and complications of fibrolamellar liver cancer through tiered computational analysis of clinical data. Nature Precision Oncology In Press.

5. Eggert, T., McGlynn, K.A., Duffy, A., Manns, M.P., Greten, T.F., and Altekruse, S.F. (2013). Fibrolamellar hepatocellular carcinoma in the USA, 2000-2010: A detailed report on frequency, treatment and outcome based on the Surveillance, Epidemiology, and End Results database. United European Gastroenterol J 1, 351–357. 10.1177/2050640613501507.

6. Dinh, T.A., Jewell, M.L., Kanke, M., Francisco, A., Sritharan, R., Turnham, R.E., Lee, S., Kastenhuber, E.R., Wauthier, E., Guy, C.D., et al. (2019). MicroRNA-375 Suppresses the Growth and Invasion of Fibrolamellar Carcinoma. Cell Mol Gastroenterol Hepatol 7, 803–817. 10.1016/j.jcmgh.2019.01.008.

7. Dinh, T.A., Utria, A.F., Barry, K.C., Ma, R., Abou-Alfa, G.K., Gordan, J.D., Jaffee, E.M., Scott, J.D., Zucman-Rossi, J., O’Neill, A.F., et al. (2022). A framework for fibrolamellar carcinoma research and clinical trials. Nat Rev Gastroenterol Hepatol. 10.1038/s41575-022-00580-3.

8. Alfarouk, K.O., Stock, C.M., Taylor, S., Walsh, M., Muddathir, A.K., Verduzco, D., Bashir, A.H., Mohammed, O.Y., Elhassan, G.O., Harguindey, S., et al. (2015). Resistance to cancer chemotherapy: failure in drug response from ADME to P-gp. Cancer Cell Int 15, 71. 10.1186/s12935-015-0221-1.

9. Cornella, H., Alsinet, C., Sayols, S., Zhang, Z., Hao, K., Cabellos, L., Hoshida, Y., Villanueva, A., Thung, S., Ward, S.C., et al. (2015). Unique genomic profile of fibrolamellar hepatocellular carcinoma. Gastroenterology 148, 806–818 e810. 10.1053/j.gastro.2014.12.028.

10. Graham, R.P., Lackner, C., Terracciano, L., Gonzalez-Cantu, Y., Maleszewski, J.J., Greipp, P.T., Simon, S.M., and Torbenson, M.S. (2018). Fibrolamellar carcinoma in the Carney complex: PRKAR1A loss instead of the classic DNAJB1-PRKACA fusion. Hepatology 68, 1441–1447. 10.1002/hep.29719.

11. Turnham, R.E., and Scott, J.D. (2016). Protein kinase A catalytic subunit isoform PRKACA; History, function and physiology. Gene 577, 101–108. 10.1016/j.gene.2015.11.052.

12. Cheung, J., Ginter, C., Cassidy, M., Franklin, M.C., Rudolph, M.J., Robine, N., Darnell, R.B., and Hendrickson, W.A. (2015). Structural insights into mis-regulation of protein kinase A in human tumors. Proc Natl Acad Sci U S A 112, 1374–1379. 10.1073/pnas.1424206112.

13. Tomasini, M.D., Wang, Y., Karamafrooz, A., Li, G., Beuming, T., Gao, J., Taylor, S.S., Veglia, G., and Simon, S.M. (2018). Conformational Landscape of the PRKACA-DNAJB1 Chimeric Kinase, the Driver for Fibrolamellar Hepatocellular Carcinoma. Sci Rep 8, 720. 10.1038/s41598-017-18956-w.

14. Turnham, R.E., Smith, F.D., Kenerson, H.L., Omar, M.H., Golkowski, M., Garcia, I., Bauer, R., Lau, H.-T., Sullivan, K.M., Langeberg, L.K., et al. (2019). An acquired scaffolding function of the DNAJ-PKAc fusion contributes to oncogenic signaling in fibrolamellar carcinoma. eLife 8, e44187. 10.7554/eLife.44187.

15. Fujimoto, A., Furuta, M., Shiraishi, Y., Gotoh, K., Kawakami, Y., Arihiro, K., Nakamura, T., Ueno, M., Ariizumi, S.-i., Hai Nguyen, H., et al. (2015). Whole-genome mutational landscape of liver cancers displaying biliary phenotype reveals hepatitis impact and molecular diversity. Nature Communications 6, 6120. 10.1038/ncomms7120.

16. 16. Wang, Z., Jensen, M.A., and Zenklusen, J.C. (2016). A Practical Guide to The Cancer Genome Atlas (TCGA). In Statistical Genomics: Methods and Protocols, E. Mathé, and S. Davis, eds. (Springer New York), pp. 111–141. 10.1007/978-1-4939-3578-9_6.

17. Vyas, M., Hechtman, J.F., Zhang, Y., Benayed, R., Yavas, A., Askan, G., Shia, J., Klimstra, D.S., and Basturk, O. (2020). DNAJB1-PRKACA fusions occur in oncocytic pancreatic and biliary neoplasms and are not specific for fibrolamellar hepatocellular carcinoma. Modern Pathology 33, 648–656. https://doi.org/10.1038/s41379-019-0398-2.

18. Langeberg, L.K., and Scott, J.D. (2015). Signalling scaffolds and local organization of cellular behaviour. Nature reviews. Molecular cell biology 16, 232–244. 10.1038/nrm3966.

19. Smith, F.D., Omar, M.H., Nygren, P.J., Soughayer, J., Hoshi, N., Lau, H.-T., Snyder, C.G., Branon, T.C., Ghosh, D., Langeberg, L.K., et al. (2018). Single nucleotide polymorphisms alter kinase anchoring and the subcellular targeting of A-kinase anchoring proteins. Proceedings of the National Academy of Sciences 115, E11465–E11474. 10.1073/pnas.1816614115.

20. Gopalan, J., Omar, M.H., Roy, A., Cruz, N.M., Falcone, J., Jones, K.N., Forbush, K.A., Himmelfarb, J., Freedman, B.S., and Scott, J.D. (2021). Targeting an anchored phosphatase-deacetylase unit restores renal ciliary homeostasis. Elife 10. 10.7554/eLife.67828.

21. Hinke, S.A., Navedo, M.F., Ulman, A., Whiting, J.L., Nygren, P.J., Tian, G., Jimenez-Caliani, A.J., Langeberg, L.K., Cirulli, V., Tengholm, A., et al. (2012). Anchored phosphatases modulate glucose homeostasis. EMBO J 31, 3991–4004. 10.1038/emboj.2012.244.

22. Aggarwal, S., Gabrovsek, L., Langeberg, L.K., Golkowski, M., Ong, S.E., Smith, F.D., and Scott, J.D. (2019). Depletion of dAKAP1-protein kinase A signaling islands from the outer mitochondrial membrane alters breast cancer cell metabolism and motility. J Biol Chem 294, 3152–3168. 10.1074/jbc.RA118.006741.

23. Efendiev, R., Bavencoffe, A., Hu, H., Zhu, M.X., and Dessauer, C.W. (2013). Scaffolding by A-kinase anchoring protein enhances functional coupling between adenylyl cyclase and TRPV1 channel. J Biol Chem 288, 3929–3937. 10.1074/jbc.M112.428144.

24. Scott, J.D., and Pawson, T. (2009). Cell signaling in space and time: where proteins come together and when they’re apart. Science 326, 1220–1224. 10.1126/science.1175668.

25. Smith, F.D., Esseltine, J.L., Nygren, P.J., Veesler, D., Byrne, D.P., Vonderach, M., Strashnov, I., Eyers, C.E., Eyers, P.A., Langeberg, L.K., and Scott, J.D. (2017). Local protein kinase A action proceeds through intact holoenzymes. Science 356, 1288–1293. 10.1126/science.aaj1669.

26. Scott, J.D., Dessauer, C.W., and Tasken, K. (2013). Creating order from chaos: cellular regulation by kinase anchoring. Annu Rev Pharmacol Toxicol 53, 187–210. 10.1146/annurev-pharmtox-011112-140204.

27. Chan, G.K.L., Maisel, S., Hwang, Y.C., Pascual, B.C., Wolber, R.R.B., Vu, P., Patra, K.C., Bouhaddou, M., Kenerson, H.L., Lim, H.C., et al. (2023). Oncogenic PKA signaling increases c-MYC protein expression through multiple targetable mechanisms. Elife 12. 10.7554/eLife.69521.

28. Qin, L., Guo, J., Zheng, Q., and Zhang, H. (2016). BAG2 structure, function and involvement in disease. Cell Mol Biol Lett 21, 18. 10.1186/s11658-016-0020-2.

29. Yang, K.M., Bae, E., Ahn, S.G., Pang, K., Park, Y., Park, J., Lee, J., Ooshima, A., Park, B., Kim, J., et al. (2017). Co-chaperone BAG2 Determines the Pro-oncogenic Role of Cathepsin B in Triple-Negative Breast Cancer Cells. Cell Rep 21, 2952–2964. 10.1016/j.celrep.2017.11.026.

30. Yue, X., Zhao, Y., Liu, J., Zhang, C., Yu, H., Wang, J., Zheng, T., Liu, L., Li, J., Feng, Z., and Hu, W. (2015). BAG2 promotes tumorigenesis through enhancing mutant p53 protein levels and function. Elife 4. 10.7554/eLife.08401.

31. Branon, T.C., Bosch, J.A., Sanchez, A.D., Udeshi, N.D., Svinkina, T., Carr, S.A., Feldman, J.L., Perrimon, N., and Ting, A.Y. (2018). Efficient proximity labeling in living cells and organisms with TurboID. Nat Biotechnol 36, 880–887. 10.1038/nbt.4201.

32. Omar, M.H., Lauer, S.M., Lau, H.T., Golkowski, M., Ong, S.E., and Scott, J.D. (2023). Proximity biotinylation to define the local environment of the protein kinase A catalytic subunit in adrenal cells. STAR Protoc 4, 101992. 10.1016/j.xpro.2022.101992.

33. 33. Szklarczyk, D., Gable, A.L., Lyon, D., Junge, A., Wyder, S., Huerta-Cepas, J., Simonovic, M., Doncheva, N.T., Morris, J.H., Bork, P., et al. (2019). STRING v11: protein-protein association networks with increased coverage, supporting functional discovery in genome-wide experimental datasets. Nucleic Acids Res 47, D607–d613. 10.1093/nar/gky1131.

34. Chen, E.Y., Tan, C.M., Kou, Y., Duan, Q., Wang, Z., Meirelles, G.V., Clark, N.R., and Ma’ayan, A. (2013). Enrichr: interactive and collaborative HTML5 gene list enrichment analysis tool. BMC Bioinformatics 14, 128. 10.1186/1471-2105-14-128.

35. Kuleshov, M.V., Jones, M.R., Rouillard, A.D., Fernandez, N.F., Duan, Q., Wang, Z., Koplev, S., Jenkins, S.L., Jagodnik, K.M., Lachmann, A., et al. (2016). Enrichr: a comprehensive gene set enrichment analysis web server 2016 update. Nucleic Acids Res 44, W90–97. 10.1093/nar/gkw377.

36. Xie, Z., Bailey, A., Kuleshov, M.V., Clarke, D.J.B., Evangelista, J.E., Jenkins, S.L., Lachmann, A., Wojciechowicz, M.L., Kropiwnicki, E., Jagodnik, K.M., et al. (2021). Gene Set Knowledge Discovery with Enrichr. Curr Protoc 1, e90. 10.1002/cpz1.90.

37. Bataller, R., and Brenner, D.A. (2005). Liver fibrosis. J Clin Invest 115, 209–218. 10.1172/JCI24282.

38. Zhang, P., Ma, X., Song, E., Chen, W., Pang, H., Ni, D., Gao, Y., Fan, Y., Ding, Q., Zhang, Y., and Zhang, X. (2013). Tubulin cofactor A functions as a novel positive regulator of ccRCC progression, invasion and metastasis. Int J Cancer 133, 2801–2811. 10.1002/ijc.28306.

39. Parker, A.L., Teo, W.S., McCarroll, J.A., and Kavallaris, M. (2017). An Emerging Role for Tubulin Isotypes in Modulating Cancer Biology and Chemotherapy Resistance. Int J Mol Sci 18. 10.3390/ijms18071434.

40. Hennessy, F., Nicoll, W.S., Zimmermann, R., Cheetham, M.E., and Blatch, G.L. (2005). Not all J domains are created equal: implications for the specificity of Hsp40-Hsp70 interactions. Protein Sci 14, 1697–1709. 10.1110/ps.051406805.

41. Penzo, M., Montanaro, L., Trere, D., and Derenzini, M. (2019). The Ribosome Biogenesis- Cancer Connection. Cells 8. 10.3390/cells8010055.

42. Zisi, A., Bartek, J., and Lindstrom, M.S. (2022). Targeting Ribosome Biogenesis in Cancer: Lessons Learned and Way Forward. Cancers (Basel) 14. 10.3390/cancers14092126.

43. Frankiw, L., Baltimore, D., and Li, G. (2019). Alternative mRNA splicing in cancer immunotherapy. Nat Rev Immunol 19, 675–687. 10.1038/s41577-019-0195-7.

44. Blijlevens, M., Li, J., and van Beusechem, V.W. (2021). Biology of the mRNA Splicing Machinery and Its Dysregulation in Cancer Providing Therapeutic Opportunities. Int J Mol Sci 22. 10.3390/ijms22105110.

45. Cowen, L., Ideker, T., Raphael, B.J., and Sharan, R. (2017). Network propagation: a universal amplifier of genetic associations. Nat Rev Genet 18, 551–562. 10.1038/nrg.2017.38.

46. Xu, Z., Page, R.C., Gomes, M.M., Kohli, E., Nix, J.C., Herr, A.B., Patterson, C., and Misra, S. (2008). Structural basis of nucleotide exchange and client binding by the Hsp70 cochaperone Bag2. Nat Struct Mol Biol 15, 1309–1317. 10.1038/nsmb.1518.

47. Arndt, V., Daniel, C., Nastainczyk, W., Alberti, S., and Hohfeld, J. (2005). BAG-2 acts as an inhibitor of the chaperone-associated ubiquitin ligase CHIP. Mol Biol Cell 16, 5891–5900. 10.1091/mbc.e05-07-0660.

48. Dai, Q., Qian, S.B., Li, H.H., McDonough, H., Borchers, C., Huang, D., Takayama, S., Younger, J.M., Ren, H.Y., Cyr, D.M., and Patterson, C. (2005). Regulation of the cytoplasmic quality control protein degradation pathway by BAG2. J Biol Chem 280, 38673–38681. 10.1074/jbc.M507986200.

49. Yoon, C.I., Ahn, S.G., Cha, Y.J., Kim, D., Bae, S.J., Lee, J.H., Ooshima, A., Yang, K.M., Park, S.H., Kim, S.J., and Jeong, J. (2021). Metastasis Risk Assessment Using BAG2 Expression by Cancer-Associated Fibroblast and Tumor Cells in Patients with Breast Cancer. Cancers (Basel) 13. 10.3390/cancers13184654.

50. Shabb, J.B. (2001). Physiological substrates of cAMP-dependent protein kinase. Chem Rev 101, 2381–2411. 10.1021/cr000236l.

51. Stokoe, D., Caudwell, B., Cohen, P.T., and Cohen, P. (1993). The substrate specificity and structure of mitogen-activated protein (MAP) kinase-activated protein kinase-2. Biochem J 296 ( *Pt 3*), 843–849. 10.1042/bj2960843.

52. Ueda, K., Kosako, H., Fukui, Y., and Hattori, S. (2004). Proteomic identification of Bcl2- associated athanogene 2 as a novel MAPK-activated protein kinase 2 substrate. J Biol Chem 279, 41815–41821. 10.1074/jbc.M406049200.

53. Huang, X., Shi, D., Zou, X., Wu, X., Huang, S., Kong, L., Yang, M., Xiao, Y., Chen, B., Chen, X., et al. (2023). BAG2 drives chemoresistance of breast cancer by exacerbating mutant p53 aggregate. Theranostics 13, 339–354. 10.7150/thno.78492.

54. Zhang, X., Zhang, J., Liu, Y., Li, J., Tan, J., and Song, Z. (2021). Bcl-2 Associated Athanogene 2 (BAG2) is Associated With Progression and Prognosis of Hepatocellular Carcinoma: A Bioinformatics-Based Analysis. Pathol Oncol Res 27, 594649. 10.3389/pore.2021.594649.

55. Takayama, S., Sato, T., Krajewski, S., Kochel, K., Irie, S., Millan, J.A., and Reed, J.C. (1995). Cloning and functional analysis of BAG-1: a novel Bcl-2-binding protein with anti-cell death activity. Cell 80, 279–284. 10.1016/0092-8674(95)90410-7.

56. Lee, J.H., Takahashi, T., Yasuhara, N., Inazawa, J., Kamada, S., and Tsujimoto, Y. (1999). Bis, a Bcl-2-binding protein that synergizes with Bcl-2 in preventing cell death. Oncogene 18, 6183–6190. 10.1038/sj.onc.1203043.

57. Shi, C.S., and Kehrl, J.H. (2019). Bcl-2 regulates pyroptosis and necroptosis by targeting BH3-like domains in GSDMD and MLKL. Cell Death Discov 5, 151. 10.1038/s41420-019-0230-2.

58. Chaitanya, G.V., Steven, A.J., and Babu, P.P. (2010). PARP-1 cleavage fragments: signatures of cell-death proteases in neurodegeneration. Cell Commun Signal 8, 31. 10.1186/1478-811X-8-31.

59. Broecker-Preuss, M., Becher-Boveleth, N., Muller, S., and Mann, K. (2016). The BH3 mimetic drug ABT-737 induces apoptosis and acts synergistically with chemotherapeutic drugs in thyroid carcinoma cells. Cancer Cell Int 16, 27. 10.1186/s12935-016-0303-8.

60. Townsend, P.A., Kozhevnikova, M.V., Cexus, O.N.F., Zamyatnin, A.A., Jr., and Soond, S.M. (2021). BH3-mimetics: recent developments in cancer therapy. J Exp Clin Cancer Res 40, 355. 10.1186/s13046-021-02157-5.

61. Tse, C., Shoemaker, A.R., Adickes, J., Anderson, M.G., Chen, J., Jin, S., Johnson, E.F., Marsh, K.C., Mitten, M.J., Nimmer, P., et al. (2008). ABT-263: a potent and orally bioavailable Bcl- 2 family inhibitor. Cancer Res 68, 3421–3428. 10.1158/0008-5472.CAN-07-5836.

62. Kandoth, C., McLellan, M.D., Vandin, F., Ye, K., Niu, B., Lu, C., Xie, M., Zhang, Q., McMichael, J.F., Wyczalkowski, M.A., et al. (2013). Mutational landscape and significance across 12 major cancer types. Nature 502, 333–339. 10.1038/nature12634.

63. Xu, L., Hazard, F.K., Zmoos, A.F., Jahchan, N., Chaib, H., Garfin, P.M., Rangaswami, A., Snyder, M.P., and Sage, J. (2015). Genomic analysis of fibrolamellar hepatocellular carcinoma. Hum Mol Genet 24, 50–63. 10.1093/hmg/ddu418.

64. Engelholm, L.H., Riaz, A., Serra, D., Dagnaes-Hansen, F., Johansen, J.V., Santoni-Rugiu, E., Hansen, S.H., Niola, F., and Frodin, M. (2017). CRISPR/Cas9 Engineering of Adult Mouse Liver Demonstrates That the Dnajb1-Prkaca Gene Fusion Is Sufficient to Induce Tumors Resembling Fibrolamellar Hepatocellular Carcinoma. Gastroenterology 153, 1662–1673 e1610. 10.1053/j.gastro.2017.09.008.

65. Oikawa, T., Wauthier, E., Dinh, T.A., Selitsky, S.R., Reyna-Neyra, A., Carpino, G., Levine, R., Cardinale, V., Klimstra, D., Gaudio, E., et al. (2015). Model of fibrolamellar hepatocellular carcinomas reveals striking enrichment in cancer stem cells. Nat Commun 6, 8070. 10.1038/ncomms9070.

66. Kastenhuber, E.R., Lalazar, G., Houlihan, S.L., Tschaharganeh, D.F., Baslan, T., Chen, C.C., Requena, D., Tian, S., Bosbach, B., Wilkinson, J.E., et al. (2017). DNAJB1-PRKACA fusion kinase interacts with beta-catenin and the liver regenerative response to drive fibrolamellar hepatocellular carcinoma. Proc Natl Acad Sci U S A 114, 13076–13084. 10.1073/pnas.1716483114.

67. Dinh, T.A., Vitucci, E.C., Wauthier, E., Graham, R.P., Pitman, W.A., Oikawa, T., Chen, M., Silva, G.O., Greene, K.G., Torbenson, M.S., et al. (2017). Comprehensive analysis of The Cancer Genome Atlas reveals a unique gene and non-coding RNA signature of fibrolamellar carcinoma. Sci Rep 7, 44653. 10.1038/srep44653.

68. Riggle, K.M., Riehle, K.J., Kenerson, H.L., Turnham, R., Homma, M.K., Kazami, M., Samelson, B., Bauer, R., McKnight, G.S., Scott, J.D., and Yeung, R.S. (2016). Enhanced cAMP-stimulated protein kinase A activity in human fibrolamellar hepatocellular carcinoma. Pediatr Res 80, 110–118. 10.1038/pr.2016.36.

69. Dinh, T.A., Sritharan, R., Smith, F.D., Francisco, A.B., Ma, R.K., Bunaciu, R.P., Kanke, M., Danko, C.G., Massa, A.P., Scott, J.D., and Sethupathy, P. (2020). Hotspots of Aberrant Enhancer Activity in Fibrolamellar Carcinoma Reveal Candidate Oncogenic Pathways and Therapeutic Vulnerabilities. Cell Rep 31, 107509. 10.1016/j.celrep.2020.03.073.

70. Tasken, K., and Aandahl, E.M. (2004). Localized effects of cAMP mediated by distinct routes of protein kinase A. Physiological reviews 84, 137–167. 10.1152/physrev.00021.2003.

71. Bauman, A.L., Soughayer, J., Nguyen, B.T., Willoughby, D., Carnegie, G.K., Wong, W., Hoshi, N., Langeberg, L.K., Cooper, D.M., Dessauer, C.W., and Scott, J.D. (2006). Dynamic regulation of cAMP synthesis through anchored PKA-adenylyl cyclase V/VI complexes. Mol. Cell 23, 925–931.

72. Omar, M.H., and Scott, J.D. (2020). AKAP Signaling Islands: Venues for Precision Pharmacology. Trends in Pharmacological Sciences 41, 933–946. https://doi.org/10.1016/j.tips.2020.09.007.

73. Smith, F.D., Reichow, S.L., Esseltine, J.L., Shi, D., Langeberg, L.K., Scott, J.D., and Gonen, T. (2013). Intrinsic disorder within an AKAP-protein kinase A complex guides local substrate phosphorylation. Elife 2, e01319. 10.7554/eLife.01319.

74. Mayer, M.P., and Bukau, B. (2005). Hsp70 chaperones: cellular functions and molecular mechanism. Cell Mol Life Sci 62, 670–684. 10.1007/s00018-004-4464-6.

75. Omar, M.H., Byrne, D.P., Jones, K.N., Lakey, T.M., Collins, K.B., Lee, K.S., Daly, L.A., Forbush, K.A., Lau, H.T., Golkowski, M., et al. (2022). Mislocalization of protein kinase A drives pathology in Cushing’s syndrome. Cell Rep 40, 111073. 10.1016/j.celrep.2022.111073.

76. Bathon, K., Weigand, I., Vanselow, J.T., Ronchi, C.L., Sbiera, S., Schlosser, A., Fassnacht, M., and Calebiro, D. (2019). Alterations in Protein Kinase A Substrate Specificity as a Potential Cause of Cushing Syndrome. Endocrinology 160, 447–459. 10.1210/en.2018-00775.

77. Lubner, J.M., Dodge-Kaqa, K.L., Carlson, C.R., Church, G.M., Chou, M.F., and Schwartz, D. (2017). Cushing’s syndrome mutant PKA(L)(205R) exhibits altered substrate specificity. FEBS Lett 591, 459–467. 10.1002/1873-3468.12562.

78. Walker, C., Wang, Y., Olivieri, C., Karamafrooz, A., Casby, J., Bathon, K., Calebiro, D., Gao, J., Bernlohr, D.A., Taylor, S.S., and Veglia, G. (2019). Cushing’s syndrome driver mutation disrupts protein kinase A allosteric network, altering both regulation and substrate specificity. Sci Adv 5, eaaw9298. 10.1126/sciadv.aaw9298.

79. Riehle, K.J., Yeh, M.M., Yu, J.J., Kenerson, H.L., Harris, W.P., Park, J.O., and Yeung, R.S. (2015). mTORC1 and FGFR1 signaling in fibrolamellar hepatocellular carcinoma. Mod Pathol 28, 103–110. 10.1038/modpathol.2014.78.

80. Lafaro, K.J., and Pawlik, T.M. (2015). Fibrolamellar hepatocellular carcinoma: current clinical perspectives. J Hepatocell Carcinoma 2, 151–157. 10.2147/JHC.S75153.

81. Takayama, S., and Reed, J.C. (2001). Molecular chaperone targeting and regulation by BAG family proteins. Nat Cell Biol 3, E237–241. 10.1038/ncb1001-e237.

82. Doong, H., Vrailas, A., and Kohn, E.C. (2002). What’s in the ’BAG’?--A functional domain analysis of the BAG-family proteins. Cancer Lett 188, 25–32. 10.1016/s0304-3835(02)00456-1.

83. Wang, H.Q., Zhang, H.Y., Hao, F.J., Meng, X., Guan, Y., and Du, Z.X. (2008). Induction of BAG2 protein during proteasome inhibitor-induced apoptosis in thyroid carcinoma cells. Br J Pharmacol 155, 655–660. 10.1038/bjp.2008.302.

84. Sun, L., Chen, G., Sun, A., Wang, Z., Huang, H., Gao, Z., Liang, W., Liu, C., and Li, K. (2020). BAG2 Promotes Proliferation and Metastasis of Gastric Cancer via ERK1/2 Signaling and Partially Regulated by miR186. Front Oncol 10, 31. 10.3389/fonc.2020.00031.

85. Yu, H., Ding, J., Zhu, H., Jing, Y., Zhou, H., Tian, H., Tang, K., Wang, G., and Wang, X. (2020). LOXL1 confers antiapoptosis and promotes gliomagenesis through stabilizing BAG2. Cell Death Differ 27, 3021–3036. 10.1038/s41418-020-0558-4.

86. Zhang, X., Dong, K., Zhang, J., Kuang, T., Luo, Y., Yu, J., Yu, J., and Wang, W. (2023). GNB1 promotes hepatocellular carcinoma progression by targeting BAG2 to activate P38/MAPK signaling. Cancer Sci 114, 2001–2013. 10.1111/cas.15741.

87. Kudaravalli, S., den Hollander, P., and Mani, S.A. (2022). Role of p38 MAP kinase in cancer stem cells and metastasis. Oncogene 41, 3177–3185. 10.1038/s41388-022-02329-3.

88. Oltvai, Z.N., Milliman, C.L., and Korsmeyer, S.J. (1993). Bcl-2 heterodimerizes in vivo with a conserved homolog, Bax, that accelerates programmed cell death. Cell 74, 609–619. 10.1016/0092-8674(93)90509-o.

89. Qian, S., Wei, Z., Yang, W., Huang, J., Yang, Y., and Wang, J. (2022). The role of BCL-2 family proteins in regulating apoptosis and cancer therapy. Front Oncol 12, 985363. 10.3389/fonc.2022.985363.

90. Song, Z., Xu, S., Song, B., and Zhang, Q. (2015). Bcl-2-associated athanogene 2 prevents the neurotoxicity of MPP+ via interaction with DJ-1. J Mol Neurosci 55, 798–802. 10.1007/s12031-014-0481-6.

91. Liang, S., Song, Z., Wu, Y., Gao, Y., Gao, M., Liu, F., Wang, F., and Zhang, Y. (2018). MicroRNA-27b Modulates Inflammatory Response and Apoptosis during Mycobacterium tuberculosis Infection. J Immunol 200, 3506–3518. 10.4049/jimmunol.1701448.

92. Hande, K.R. (1998). Etoposide: four decades of development of a topoisomerase II inhibitor. Eur J Cancer 34, 1514–1521. 10.1016/s0959-8049(98)00228-7.

93. Potluri, J., Harb, J., Masud, A.A., and Huu, J.E. (2020). A Phase 3, Double-Blind, Placebo-Controlled, Randomized Study Evaluating Navitoclax in Combination with Ruxolitinib in Patients with Myelofibrosis (TRANSFORM-1). Blood 136, 4–4. 10.1182/blood-2020-139758.

94. Harrison, C.N., Garcia, J.S., Somervaille, T.C.P., Foran, J.M., Verstovsek, S., Jamieson, C., Mesa, R., Ritchie, E.K., Tantravahi, S.K., Vachhani, P., et al. (2022). Addition of Navitoclax to Ongoing Ruxolitinib Therapy for Patients With Myelofibrosis With Progression or Suboptimal Response: Phase II Safety and Efficacy. J Clin Oncol 40, 1671–1680. 10.1200/JCO.21.02188.

