## Supplemental Figures for "Recruitment of BAG2 to DNAJ-PKAc scaffolds promotes cell survival and resistance to drug-induced apoptosis in fibrolamellar carcinoma"

**<PHOTOACTIVATION VIDEOS>**

**Supplemental Figure 2.**

Video S1. WT PKAc remains localized within AKAP-signaling islands, related to Figure 2. AML12 hepatocytes expressing AKAP79-YFP (cyan), RIIα-iRFP, and PKAc-WT tagged with photoactivatable mCherry (magenta). Movie shows 2 seconds of baseline fluorescence prior to photoactivation and 13 seconds of recording after with sampling every 500 ms.

Video S2. DNAJ-PKAc diffuses into the cytosol away from AKAP-signaling islands, related to Figure 2. AML12 hepatocytes expressing AKAP79-YFP (cyan), RIIα-iRFP, and DNAJ-PKAc tagged with photoactivatable mCherry (magenta). Movie shows 2 seconds of baseline fluorescence prior to photoactivation and 13 seconds of recording after with sampling every 500 ms.

Video S3. DNAJ$^{H33Q}$-PKAc demonstrates an intermediate phenotype, related to Figure 2. AML12 hepatocytes expressing AKAP79-YFP (cyan), RIIα-iRFP, and DNAJ$^{H33Q}$-PKAc tagged with photoactivatable mCherry (magenta). Movie shows 2 seconds of baseline fluorescence prior to photoactivation and 13 seconds of recording after with sampling every 500 ms.

Video S4. PKAc$^{\Delta 14}$ remains localized within AKAP-signaling islands, related to Figure 2. AML12 hepatocytes expressing AKAP79-YFP (cyan), RIIα-iRFP, and PKAc $^{\Delta 14}$ tagged with photoactivatable mCherry (magenta). Movie shows 2 seconds of baseline fluorescence prior to photoactivation and 13 seconds of recording after with sampling every 500 ms.

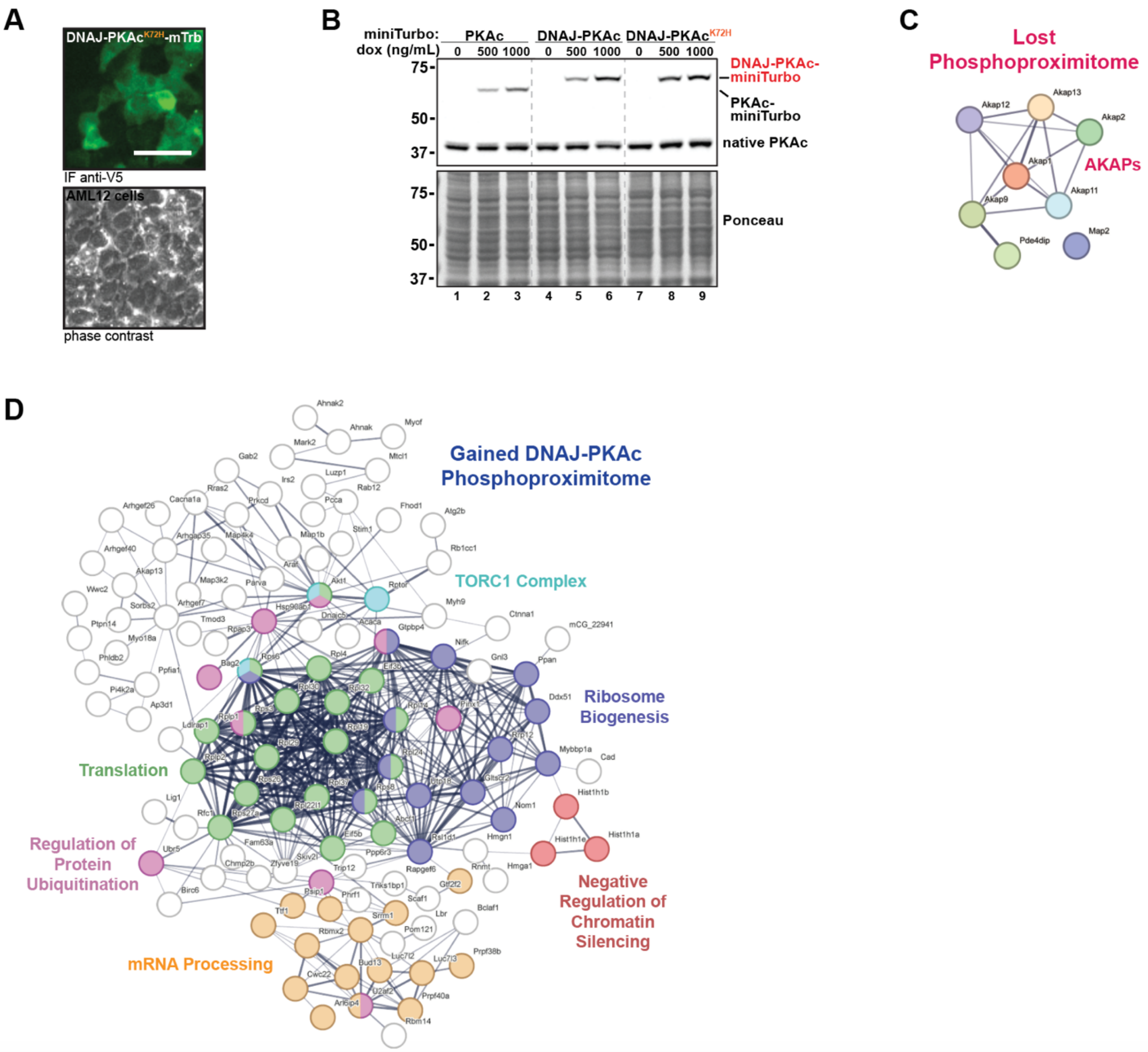
A
DNAJ-PKAc^K72H-mTrb
IF anti-V5
AML12 cells
phase contrast
B
miniTurbo:
dox (ng/mL)
PKAc
DNAJ-PKAc
DNAJ-PKAc^K72H
0
500
1000
75
50
37
DNAJ-PKAc-miniTurbo
PKAc-miniTurbo
native PKAc
Ponceau
1
2
3
4
5
6
7
8
9
C
Lost Phosphoproximitome
Akap12
Akap13
Akap2
Akap1
Akap11
Akap9
Pde4dip
Map2
AKAPs
D
Gained DNAJ-PKAc Phosphoproximitome
TORC1 Complex
Ribosome Biogenesis
Translation
Regulation of Protein Ubiquitination
Negative Regulation of Chromatin Silencing
mRNA Processing

E

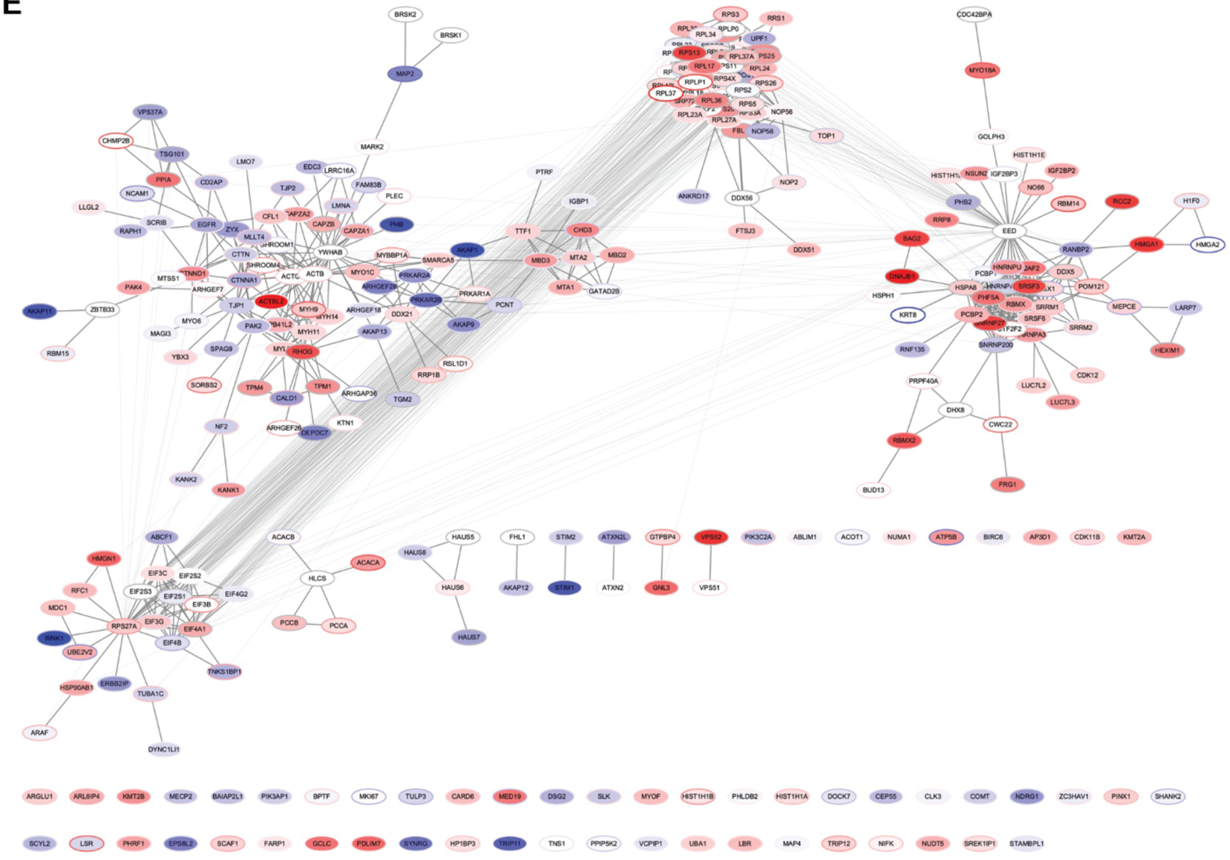


**Supplemental Figure 3.**

A) Immunofluorescence imaging of AML12 hepatocytes demonstrating inducible expression of DNAJ-PKAc$^{K72H}$-mTrb (green, top) with corresponding phase contrast (bottom). Scale bar = 50 μm.

B) Immunoblot of cell lysate from AML12 stable lines demonstrating doxycycline-inducible expression of miniTurbo-tagged PKAc variants (top bands) and native PKAc (bottom bands).

C) STRING network depiction of selected phosphoproteins with lesser enrichment in DNAJ-PKAc versus WT PKAc.

D) STRING network depiction of selected phosphoproteins with greater enrichment in DNAJ-PKAc versus WT PKAc. Node color corresponds with similarly colored annotation.

E) Depiction of results from ReactomeFI network propagation. Proximity proteomics is respresented by node fill color. Proximity phosphoproteomics is represented by node border color. Degree of association is represented on a yellow (more associated) to blue (less associated) spectrum based on $\log_2$(fold change). FDR < 0.25.

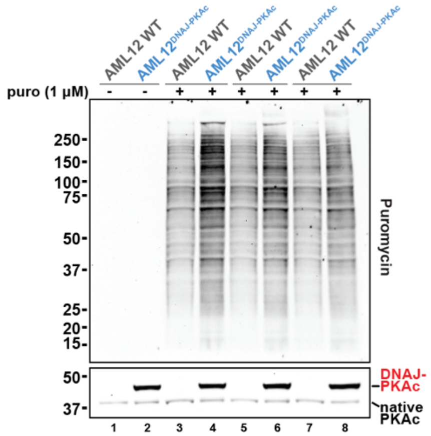


**Supplemental Figure 4.**
Full immunoblot showing three biological replicates of cell lysates from WT AML12 and AML12$^{DNAJ\text{-}PKAc}$ treated with either vehicle or puromycin (50 µM). Puromycin in top panel shows newly synthesized, puromycin-labeled proteins. PKAc in bottom panel shows expression of miniTurbo-tagged PKAc variants (top band) over native PKAc (bottom band).

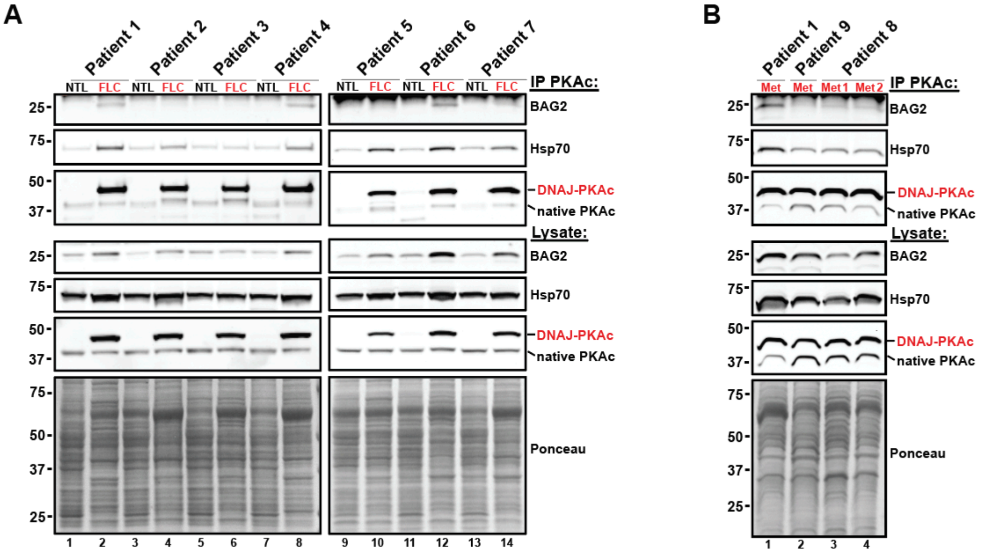


**Supplemental Figure 5.**

A) Immunoprecipitation with antibody against PKAc from paired normal adjacent liver (NTL) and FLC tumor (FLC) tissue lysates from 7 patients.
B) Immunoprecipitation with antibody against PKAc from tumor tissue lysates of four metastatic (Met) recurrences across 3 patients.
